# Secondary metabolism in the gill microbiota of shipworms (Teredinidae) as revealed by comparison of metagenomes and nearly complete symbiont genomes

**DOI:** 10.1101/826933

**Authors:** Marvin A. Altamia, Zhenjian Lin, Amaro E. Trindade-Silva, Iris Diana Uy, J. Reuben Shipway, Diego Veras Wilke, Gisela P. Concepcion, Daniel L. Distel, Eric W. Schmidt, Margo G. Haygood

## Abstract

Shipworms play critical roles in recycling wood in the sea. Symbiotic bacteria supply enzymes that the organisms need for nutrition and wood degradation. Some of these bacteria have been grown in pure culture and have the capacity to make many secondary metabolites. However, little is known about whether such secondary metabolite pathways are represented in the symbiont communities within their hosts. In addition, little has been reported about the patterns of host-symbiont co-occurrence. Here, we collected shipworms from the United States, the Philippines, and Brazil, and cultivated symbiotic bacteria from their gills. We analyzed sequences from 22 shipworm gill metagenomes from seven shipworm species and from 23 cultivated symbiont isolates. Using (meta)genome sequencing, we demonstrate that the cultivated isolates represent all the major bacterial symbiont species and strains in shipworm gills. We show that the bacterial symbionts are distributed among shipworm hosts in consistent, predictable patterns. The symbiotic bacteria encode many biosynthetic gene cluster families (GCFs) for bioactive secondary metabolites, only <5% of which match previously described biosynthetic pathways. Because we were able to cultivate the symbionts, and sequence their genomes, we can definitively enumerate the biosynthetic pathways in these symbiont communities, showing that ∼150 out of ∼200 total biosynthetic gene clusters (BGCs) present in the animal gill metagenomes are represented in our culture collection. Shipworm symbionts occur in suites that differ predictably across a wide taxonomic and geographic range of host species, and collectively constitute an immense resource for the discovery of new biosynthetic pathways to bioactive secondary metabolites.

**Importance:** We define a system in which the major symbionts that are important to host biology and to the production of secondary metabolites can be cultivated. We show that symbiotic bacteria that are critical to host nutrition and lifestyle also have an immense capacity to produce a multitude of diverse and likely novel bioactive secondary metabolites that could lead to the discovery of drugs, and that these pathways are found within shipworm gills. We propose that, by shaping associated microbial communities within the host, the compounds support the ability of shipworms to degrade wood in marine environments. Because these symbionts can be cultivated and genetically manipulated, they provide a powerful model for understanding how secondary metabolism impacts microbial symbiosis.

## Introduction

Shipworms (Family Teredinidae) are bivalve mollusks found throughout the world’s oceans (1, 2). Many shipworms eat wood, assisted by cellulases from intracellular symbiotic γ- proteobacteria that inhabit their gills (Fig. 1) (3–6). Other shipworms use sulfide metabolism, also relying on gill-dwelling γ-proteobacteria for sulfur oxidation (7). Shipworm gill symbionts of several different species are thus essential to shipworm nutrition and survival. One of the most remarkable features of the shipworm system is that wood digestion does not take place where the bacteria are located, so that the bacterial cellulase products are transferred from the gill to a nearly sterile cecum (8), where wood digestion occurs (Fig. 1) (9). This enables the host shipworms to directly consume glucose and other sugars derived from wood lignocellulose and hemicellulose, rather than the less energetic fermentation byproducts of cellulolytic gut microbes as found in other symbioses. Shipworm symbionts are also essential for nitrogen fixation that helps to offset the low nitrogen content of wood (10, 11). Thus, shipworms have evolved structures and mechanisms enabling bacterial metabolism to support animal host nutrition.

**Figure 1.**
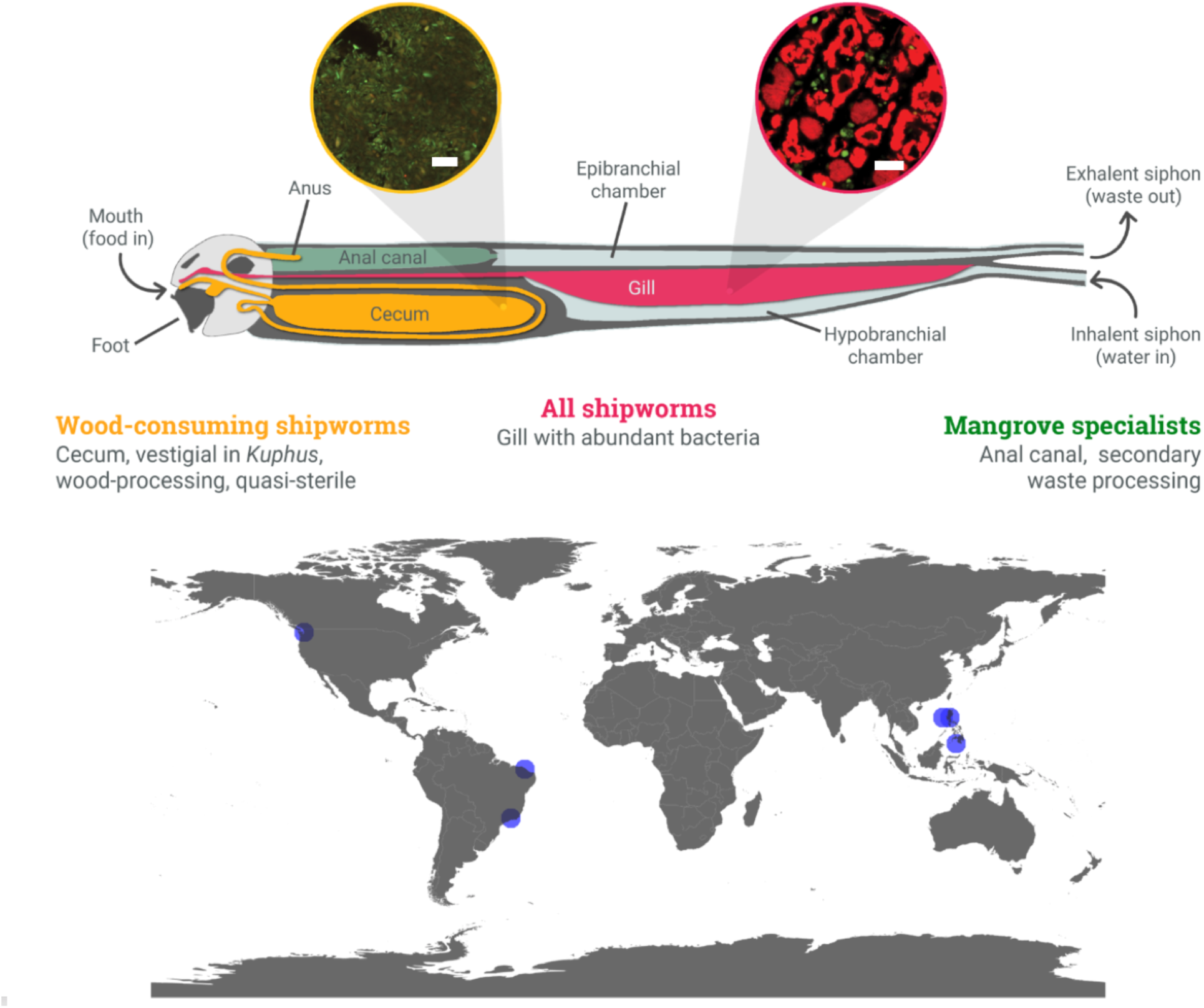
Top, diagram of generic shipworm anatomy. Insets are from Betcher *et al*., *PLoS One*, 2012 Figure 2, panels B and D, scale bar 20 µm (8). Red: signal from a fluorescent universal bacterial probe indicating large numbers of bacterial symbionts in the bacteriocytes of the gill, and paucity of bacteria in the cecum. Green is background fluorescence. Bottom, collection locations of specimens included in this study. See Table S1 for details.

While in many nutritional symbioses the bacteria are difficult to cultivate, shipworm gill symbiotic γ-proteobacteria have been brought into stable culture (5, 12, 13). This led to the discovery that these bacteria are exceptional sources of secondary metabolites (14). Of bacteria with sequenced genomes, the gill symbionts *Teredinibacter turnerae* T7901 and related strains are among the richest sources of biosynthetic gene clusters (BGCs), comparable in content to famous producers of commercial importance such as *Streptomyces* spp. (13–16). This implies that shipworms might be a good source of new compounds for drug discovery. Of equal importance, the symbiotic bacteria are crucial to survival of host shipworms, and bioactive secondary metabolites might play a role in shaping those symbioses.

An early analysis of the *turnerae* T7901 genome revealed nine complex polyketide synthase (PKS) and nonribosomal peptide synthetase (NRPS) BGCs (14). One of these was shown to produce a novel catecholate siderophore, turnerbactin, which is crucial in obtaining iron and to the survival of the symbiont in nature (17). A second BGC synthesizes the borated polyketide tartrolons D/E, which are antibiotic and potently antiparasitic compounds (18). Both were detected in the extracts of shipworms, implying a potential role in producing the remarkable near sterility observed in the cecum (8). These data suggested specific roles for secondary metabolism in shipworm ecology.

*T. turnerae* T7901 is just one of multiple strains and species of γ-proteobacteria living intracellularly in shipworm gills (3, 12), and thus these analyses just begin to describe shipworm secondary metabolism. Many shipworm species are generalists, consuming wood from a variety of sources (1, 19). Other wood-eaters, such as *Dicyathifer mannii*, *Bactronophorus thoracites*, and *Neoteredo reynei*, are specialists that live in the submerged branches, trunks and rhizomes of mangroves (20, 21). There, they play an important role in ecological processes in mangrove ecosystems, i.e. transferring large amount of carbon fixed by mangroves to the marine environment (19). Several shipworm species, such as *Kuphus polythalamius*, live in other substrates. *K. polythalamius* often is found in sediment habitats (as well as in wood) where its gill symbionts are crucial to sulfide oxidation and carry out carbon fixation (7). *K. polythalamius* lacks significant amounts of cellulolytic symbionts such as *T. turnerae*, and instead contains *Thiosocius teredinicola*, which oxidizes sulfide and generates energy for the host (22). Other shipworms are found in solid rock and in seagrass (23, 24). Thus, gill symbionts vary, but in all cases the symbionts appear to be essential to the survival of shipworms.

While the potential of *T. turnerae* as an unexplored producer of secondary metabolites has been described (14, 16), the capacity of other shipworm symbionts is still largely unknown. Moreover, several facts indicate that the BGCs found in cultivated isolates might also be produced by symbionts within shipworm gills, but their presence, distribution and variability in nature are unknown. Previous data include the detection of tartrolons and turnerbactins and their BGCs in shipworms (17, 18); an investigation of four isolate genomes and one metagenome that observed shared pathways (25); also an exploratory investigation of the metagenome of *N. reynei* gills and digestive tract led to the detection of known *T. turnerae* BGCs as well as novel clusters (26). These findings left major questions about the origin, abundance, variability, distribution, and potential roles of shipworm secondary metabolites.

Here, we use a comparative metagenomics approach to answer these questions. We selected six species of wood-eating shipworms (*B. thoracites*, *N. reynei*, *Bankia setacea*, *Bankia* sp., D*. mannii*, and *Teredo* sp.), comparing these to a seventh sulfide-oxidizing group, *Kuphus* spp. We compared gill metagenomes from 22 specimens comprising seven animal species with the genomes of 23 cultivated bacteria isolated from shipworms. These isolated bacteria included 22 cellulolytic and sulfur-oxidizing isolates cultivated from shipworm tissue samples. By comparing the gill metagenomes to isolate strain genomes, we demonstrate that the cultivated bacterial genomes accurately represent the genomes of symbionts found in the gills, and we show that they share many of the same secondary metabolic BGCs. Moreover, we show that the members of symbiont communities differ among shipworm species, indicating that surveying more host shipworms will lead to discovery of new BGCs and new bacterial symbionts.

## Results and Discussion

### Sequencing data

Most of the genomes and metagenomes were obtained in this work and are described here for the first time, or in a few cases previously reported genomes/metagenomes were resequenced/reassembled/reanalyzed (see Methods). Two bacterial genomes, *T. turnerae* T7901 and *T. teredinicola* 2141T, and metagenomes of *K. polythalamius* were previously described (7, 14, 22). The resulting statistics and accession numbers are provided in Table S1 A, while specimen and strain origin, many of which have not been previously reported, are given in Table S1 B. For bacterial strains, six of the circular genomes were closed, while remaining assemblies had between 2-141 scaffolds. Metagenome total assembly sizes ranged from 2.6×10^8^-1.3×10^9^ bp, with N50s of 860-4530 bp. The larger N50s were obtained with the Philippines specimens sequenced at the University of Utah, while others sequenced elsewhere had comparatively shorter N50s.

### Mapping cultivated bacteria to gill metagenomes

A phylogenetic tree created from the 16S rRNA genes of the cultivated bacteria (Fig. 2A and S1) revealed that the strains are all γ- Proteobacteria. Of these, 21 are from Order *Cellvibrionales,* including 11 strains of *T. turnerae*, and 10 strains of diverse cellulolytic bacteria, most of which have not been previously described. An exception is strain BS02, which was recently formally described as the new species, *Teredinibacter waterburyi* (27). The remaining two strains are from Order *Chromatiales* (*Thiosocius* and allies).

**Figure 2.**
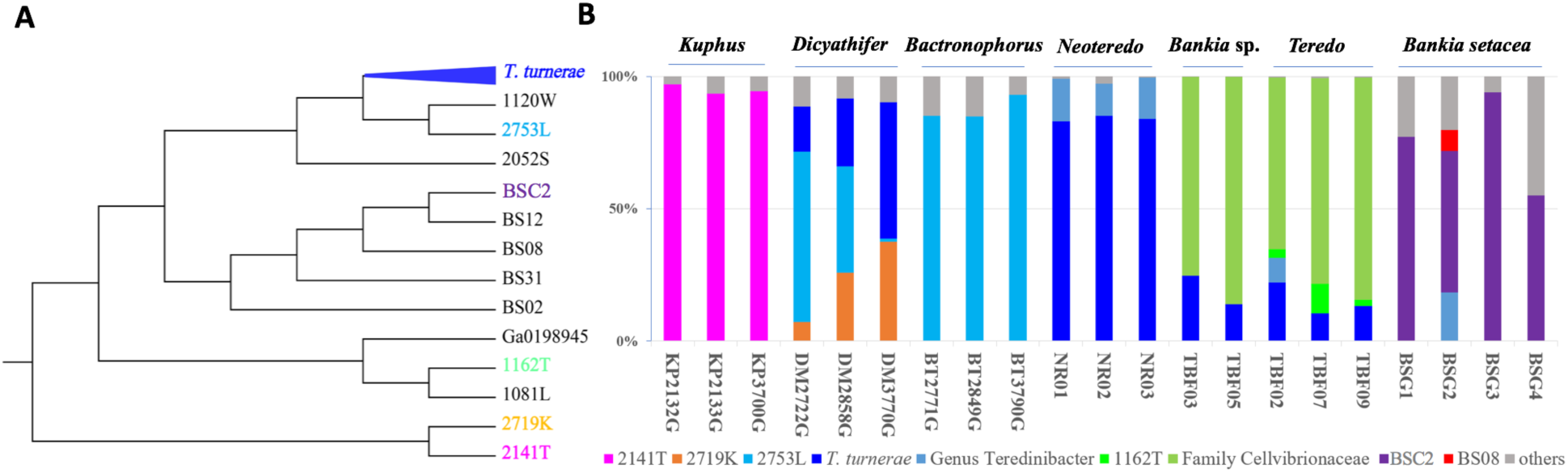
Cultivated bacterial isolates represent the major shipworm gill symbionts. **A)** Isolated bacteria analyzed in this study are shown in abstracted schematic of a 16S rRNA phylogenetic tree. The complete tree with accurate branch lengths and bootstrap numbers is shown in Fig. S1. *T. turnerae* comprised 11 sequenced strains, for other groups individual strains are shown. Each color indicates different bacteria appearing in the metagenomes in B. **B)** Species composition of shipworm gill symbiont community based on shotgun metagenome sequence analysis. The *y*-axis indicates the percent of reads originating from each bacterial species, while the *x*-axis indicates individual shipworm specimens used in the study. Colors indicate the origin of bacterial reads; gray is minor, sporadic, unidentified strains.

**Figure 3.**
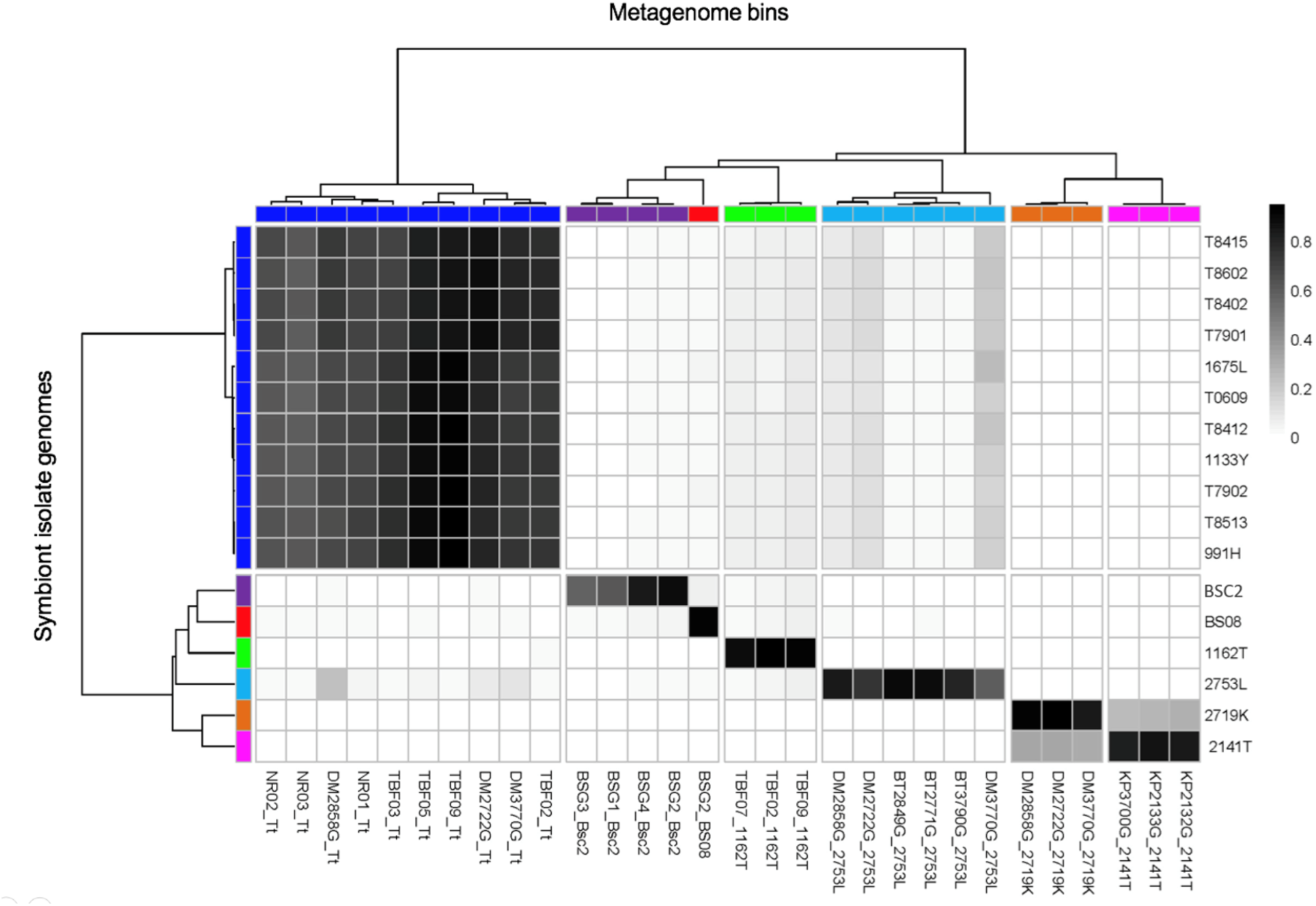
Heatmap of relationships between symbiont isolate genomes and gill metagenome bins. The scale bar is shaded according to identity based upon (AF x gANI). Color bars in the phylogenetic tree indicate bacterial species identity, either in the metagenomes or in the genome, and they are identical to the codes shown in Fig. 2. This figure indicates the high degree of certainty that the cultivated isolates are the same species as the major bacteria present in the gill.

Further, gANI measurements reinforce the 16S rRNA based phylogenetic tree of sequenced strains (Fig. 2, 3, and S1, Table S2). Previously proposed cut-offs for bacterial species differentiation suggest that bacterial strains with gANI values ≥0.95 are conspecific, although several well-known species have lower gANI values (28). The concatenated *T. turnerae* strains are represented by two groups, exemplified by strains T7901 and T7902 (Fig. S2). Within each group, *T. turnerae* strains have gANI values >0.97, whereas between groups the gANI values are ∼0.92. This agrees with and reinforces a previously published observation that *T. turnerae* is comprised of two distinct clades and suggests that these clades may in fact constitute distinct but closely related bacterial species (12). Outside of *T. turnerae*, the strains are much less closely related, with AF x gANI values <0.4 (Figs. 3 and S2), indicating that they are all different at the species level.

Using metagenomic methods, the bacteria living in gills were grouped into bins that represent individual species of bacteria (Fig. 2B). For example, in *Kuphus* spp., >95% of bacterial reads could be mapped to cultivated isolate strain *T. teredinicola* 2141T. Among the three specimens measured, 14 bins mapped to *T. teredinicola* 2141T (Table S2). None of the other specimens in our study had any match to *T. teredinicola* 2141T with gANI >0.90. Normalized by length, these bins had a total gANI = 0.96 (Table 1). In comparison to values obtained in the phylogenetic tree, these data suggest that *T. teredinicola* 2141T is conspecific with the uncultivated symbionts in the metagenomes of *Kuphus* spp.

**Table 1.** Example gANI values for shipworm gills in comparison to sequenced isolates, extracted from Table S2.

| Comparison | Total metagenomic bin size in bp | gANI |
| --- | --- | --- |
| *Kuphus* spp. : *T. teredincicola* 2141T | 23758169 | 0.963839 |
| *D. mannii* and *B. thoracites* : 2753L | 26758239 | 0.990273 |
| *D. mannii* : 2719K | 14895102 | 0.992012 |
| *B. setacea* : BSC2 | 19687153 | 0.974108 |
| *Teredo* sp. : 1162T | 1489154 | 0.984036 |
| *B. setacea* : BS08 | 3513639 | 0.995137 |

Similarly, *Cellvibrionaceae* strain 2753L was mapped to 20 bins in *D. mannii* and *B. thoracites* specimens, with a total gANI >0.99. When bins were mapped to discrete strains as shown in Fig. 2B, the gANI was 0.96-0.99 to a single strain, with much lower identity to other strains sequenced. These data demonstrate a high level of identity between cultivated isolates and the strains present within shipworm gills, suggesting that in some cases these are near identical strains to those present within the shipworms.

In other cases, either because we had multiple strains representing a species (as in *T. turnerae*) or because the identity to single strains was not as pronounced, we described bins as “*T. turnerae*”, “Genus Teredinibacter”, and “Family Cellvibrionaceae”. These still had relatively high identities to cultivated isolates. For example, the *Teredo* sp. bins in total had a gANI > 0.98 to cultivated isolates in our strain collection. It is likely that the metagenomes from these animals were not as similar to cultivated isolates because, in those cases, we compared isolates from Philippines specimens with metagenomes of Brazilian animals.

In sum, these data demonstrate conclusively that the cultivated isolates obtained from shipworm gills accurately represent the strains found within shipworms. The data suggest that the isolates are the same species as the true symbionts found within the animals, and in many cases they are >99% DNA sequence identical at the whole genome level. The data reveal that >85% of the DNA in each specimen’s gill metagenome is represented by a cultivated isolate in our collection (with the exception of one specimen), and that the remaining <15% of the DNA belongs to multiple, low-abundance species, most of which are not reproducibly found in multiple shipworm isolates. Further, much of the shipworm literature focuses on the readily cultivable *T. turnerae*. We show that *T. turnerae* is dominant in some species, but it is very minor or even absent in others.

### Strain variation increases genetic diversity of shipworm microbiota

Metagenome binning defined the major symbiont species present in shipworm gills to be a relatively simple mixture of one to three species. Since we had deep sequencing of the major metagenomic bacterial species, we expected to be able to provide complete assemblies. In other instances, using similarly deep data, we have been able to obtain relatively complete assemblies, or even to assemble whole bacterial genomes from metagenomes (29). However, our metagenome bin N50s were only in the very low thousands.

When investigating the causes underlying the challenge of assembly, we noted that we often obtained very similar contigs with different copy numbers. For example, a single metagenome bin containing BCS2-like contigs is shown in Table S3. Pairwise identities between contigs are very high, between 93-98% DNA sequence identical, indicating that these bins are comprised of mixtures of very closely related bacteria. We saw a very similar phenomenon in a recent investigation of *K. polythalamius* symbionts (7). In that case, the strains were nearly identical and could not be resolved by 16S rRNA gene sequences, which were 100% identical. Thus, we developed a different method to quantify strain-level variation that was observed using metagenomics.

In the *Kuphus* study, we cut the DNA gyrase B gene into 50 bp segments and aligned single reads to each 50 bp segment (7). By quantifying reads for each observed SNP, we confirmed that the gill symbiont species consisted of several strains, and we quantified their relative abundances. Here, we expanded this previous knowledge by investigating the major strains found in the remaining shipworm species, using the same method. We show four additional examples (Fig. S3) in which we can quantify multiple strains of each different bacterial symbiont species, but the same phenomenon obtains in all of the metagenomes. This analysis shows that similar strain variation is a widespread phenomenon in shipworm gills, and not just restricted to *K. polythalamius*. We believe that strain variation is likely to be an important source of BGC variation, as described further below.

### Discovery and analysis of BGCs

Knowing that the cultivated bacteria represent the major symbiont species present in gill metagenomes, we next compared secondary metabolism between these specimens and isolates. To start, we took an inventory of the BGC content in our assembled sequences. Analysis using antiSMASH (30) revealed a large number of BGCs: 431 BGCs were identified in the 23 cultivated isolates alone. Because raw antiSMASH output includes many hypothetical or poorly characterized BGCs, we chose to focus on well-characterized classes of secondary metabolic proteins and pathways: polyketide synthases (PKSs), nonribosomal peptide synthetases (NRPSs), siderophores, terpenes, homoserine lactones, and thiopeptides. Using these criteria, we identified 168 BGCs from 23 cultivated isolates and 401 BGCs from 22 shipworm gill metagenomes (Fig. 4). Because the genomes of cultivated isolates were well assembled, we could discern and analyze entire BGCs. By contrast, animal metagenomes had smaller contigs, so that BGCs were fragmented.

**Figure 4.**
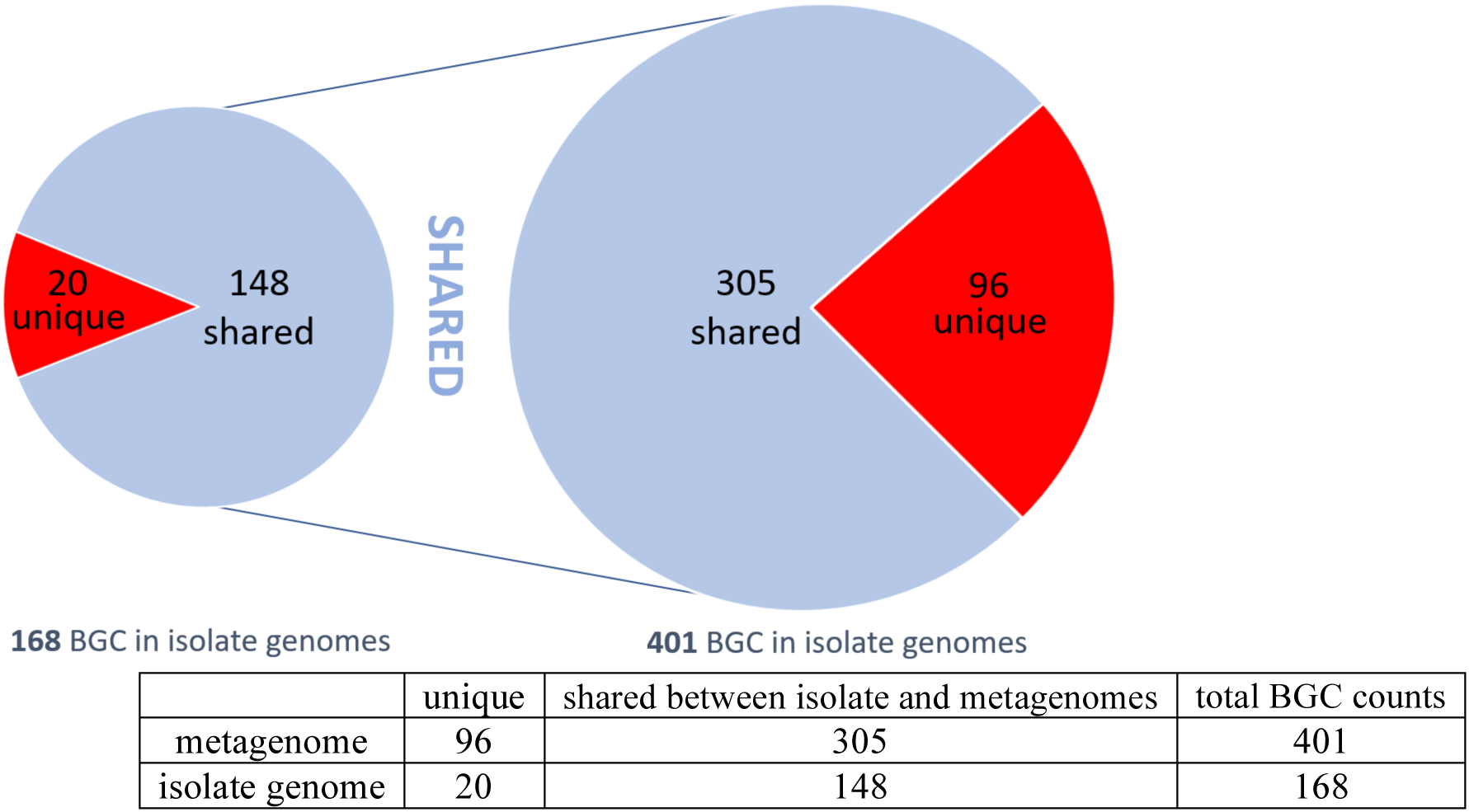
Most BGCs found in the metagenomes and in the bacterial isolate genomes are shared. 401 BGCs from metagenome sequences were compared to the bacterial isolate genomes, of which 305 could be found in isolates. Conversely, 148 of 168 BGCs from sequenced bacterial isolates could be found in the metagenomes. The shared numbers likely differ because the contigs assembled from the metagenome sequences were shorter on average, so that several metagenome fragments may map to a single BGC in an isolate.

The BGCs identified in this study nearly universally originate from Order *Cellvibrionales*, with very few BGCs found in the sulfide oxidizing strains *Chromatiales*. Thus, the cellulolytic shipworm symbionts are rich sources of diverse BGCs. We found only five BGCs that were similar to previously identified clusters from outside of shipworms, based upon >70% of genes conserved in antiSMASH. The remainder appeared to be unknown or uncharacterized BGCs. In turn, the new BGCs are likely to represent new compounds, while characterized BGCs represent those for previously identified compounds. In addition, it is possible that some of the new BGCs may represent known compounds, for which biosynthetic pathways have not yet been discovered. This result further supports a previous analysis comparing genomes across domain Bacteria, which revealed that *T. turnerae* represents a notably rich, yet nearly untapped, source of new secondary metabolite genes (16).

To facilitate comparison between metagenomes, we grouped all 569 BGCs into 122 gene cluster families (GCFs), where each GCF is comprised of closely related BGCs (31, 32) (Fig. 5 and Table S4). BGCs grouped into a single GCF are highly likely to encode the production of identical or closely related secondary metabolites.

**Figure 5.**
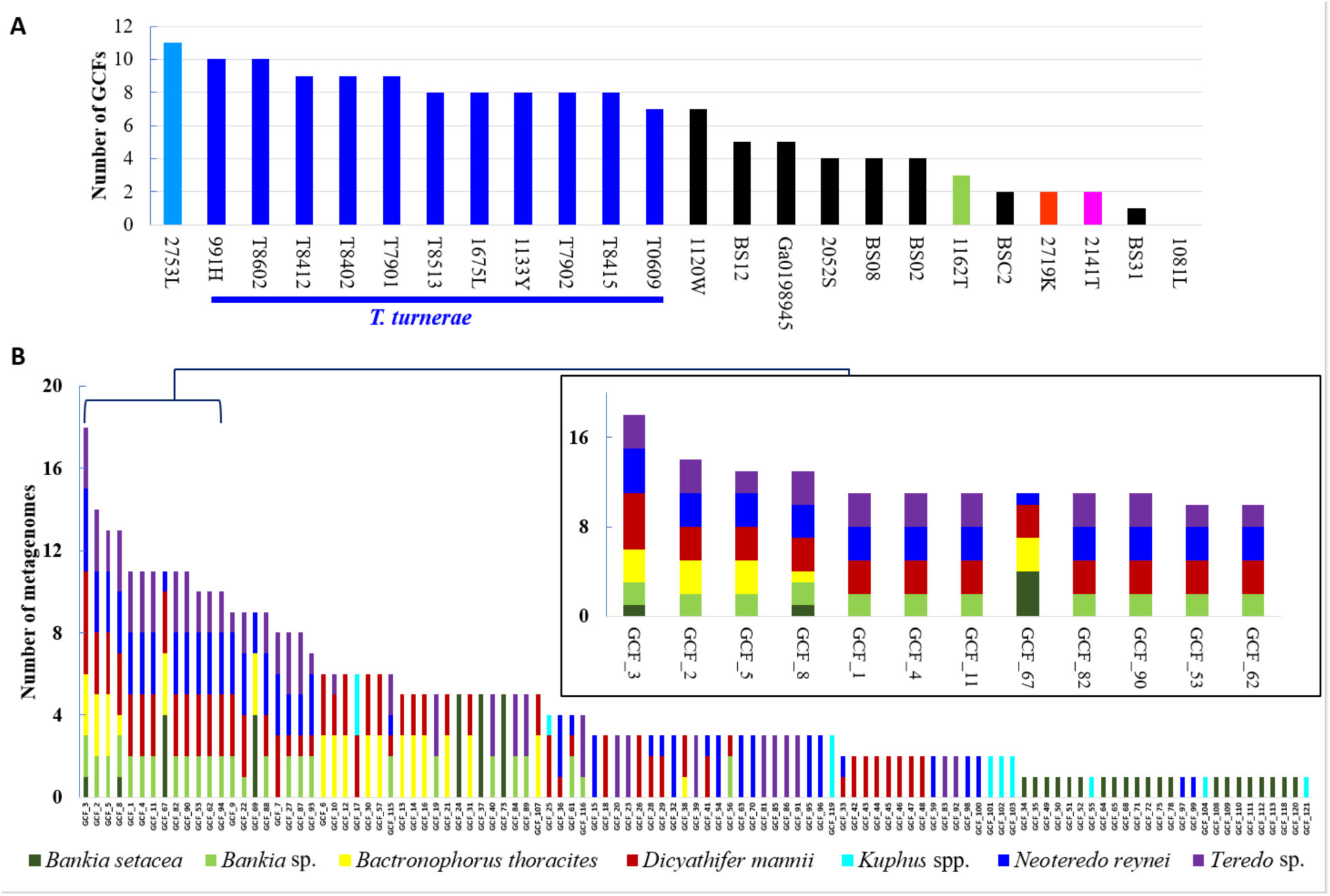
GCFs found in **A)** bacterial genomes and **B)** gill metagenomes. **A)** A list of strains of cultivated bacterial genomes is provided in the *x*-axis, while the number of total GCFs in different sequenced strains is shown in the *y*-axis. Colors indicate bacteria from Fig. 2A. Because there are 11 isolates of a *T. turnerae*, the number of GCFs in this group (dark blue bars) are comparatively overrepresented in the diagram. **B)** GCFs (*x*-axis) found in each metagenome (*y*-axis) are shown. The inset expands a region containing the most common GCFs found in our specimens. Colors indicate shipworm host species. See Table S4 for a complete list of GCFs used in this figure.

Some important BGCs were excluded using our method. For example, we analyzed the genome of *Chromatiales* strain 2719K and discovered a gene cluster for tabtoxin (33, 34) or a related compound (Fig. 6). This cluster does not contain common PKS/NRPS elements and thus is not one of the GCFs shown in Figures 5, 7, or 8. A key biosynthetic gene in the tabtoxin-like cluster was pseudogenous in strain 2719K, but the *D. mannii* gill metagenome contained an apparently functional pathway. Tabtoxin is an important β-lactam that is used by *Pseudomonas* in plant pathogenesis (35, 36).

**Figure 6.**
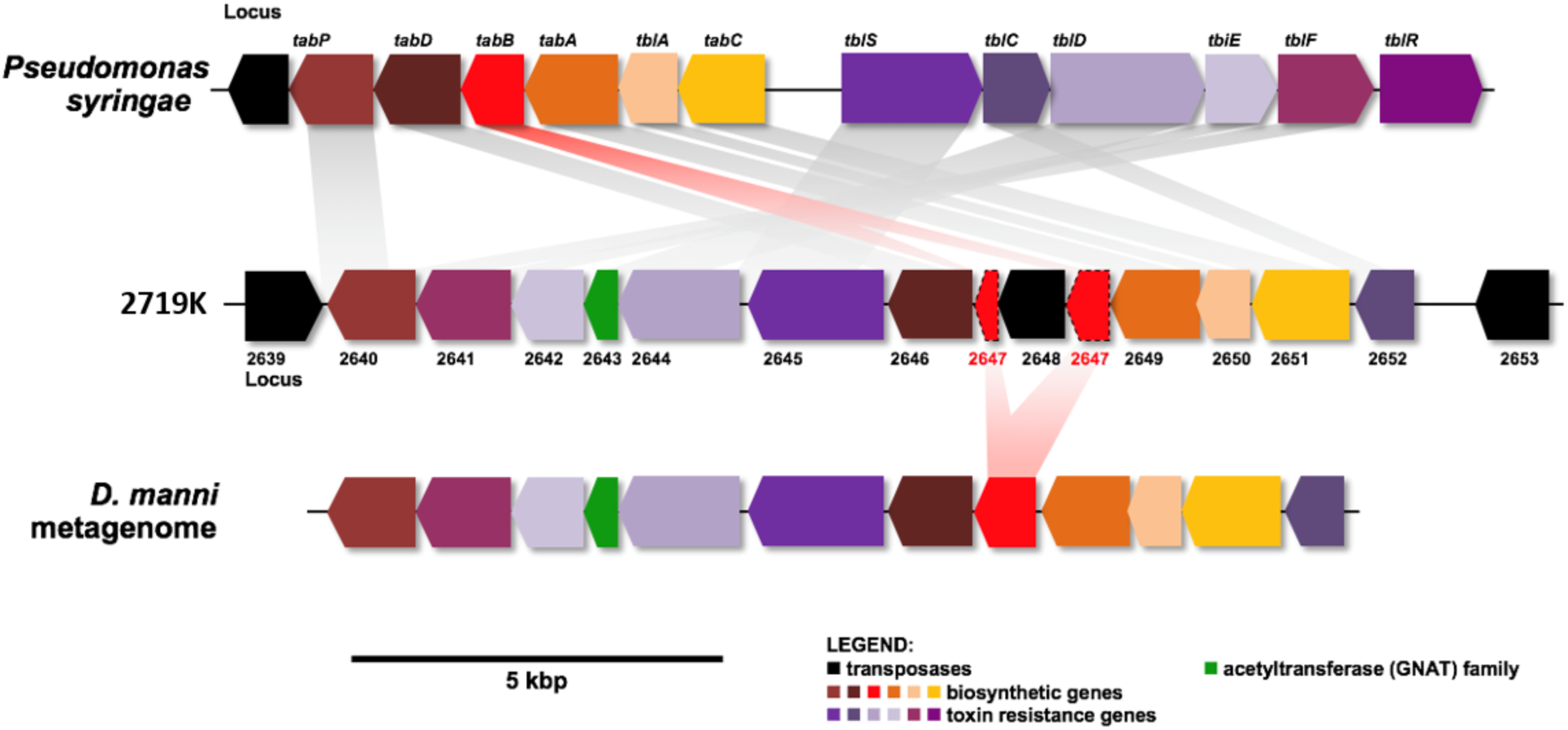
A possible tabtoxin pathway is found in the *D. manni* metagenome. Tabtoxin is a phytotoxin β-lactam initially discovered in *Pseudomonas* spp. (top). Strain 2719K contained a tabtoxin-like cluster that was pseudogenized (shown as an insertion in *tabB*; middle). A non-pseudogenized tabtoxin-like cluster was found in the *D. manni* metagenome gill (bottom) supporting the observation that multiple variants of each symbiont genome are represented in each metagenome.

### Comparison of isolate and gill BGCs

Of 401 BGCs identified in the metagenomes, 305 of them also had close relatives in cultivated isolates, indicating that ∼75% of BGCs in the metagenomes are covered in our sequenced culture collection (Fig. 4). Conversely, of 168 isolate BGCs, 148 (90%) of them are found in the metagenomes. Thus, sequencing additional cultivated isolates in our strain collections is likely to yield additional novel BGCs. Since the 11 *T. turnerae* strains analyzed in this project contain different BGCs, we speculate that the additional BGC variation is due to the observed strain variation in the shipworm gills.

It is notoriously difficult to quantify BGCs in metagenomes, which usually contain relatively small contigs. Since BGCs in the classes that we analyzed are usually between 10 kbp to >100 kbp in length, each BGC is usually represented by multiple, short contigs, which are not easily mapped. Here, we had an advantage in that the cultivated isolates accurately represented the gill metagenomes: we could map the identified metagenomic contigs to the assembled BGCs found in cultivated isolates.

Using this mapping, we could accurately estimate the number of unique BGCs in the gill symbiont community. For example, 305 metagenome BGCs are synonymous with 148 isolate BGCs, indicating that the metagenome BGC count can be estimated to be approximately double the actual number of BGCs. To verify this estimate, we selected GCFs 2, 3, 5 and 8, aligning metagenomic contigs against the BGCs from cultivated isolates (Fig. S4). In the metagenomes, out of the 401 total BGCs identified, 100 were members of these four GCFs, but some of them were just fragments of the full-length BGCs found in cultivated isolates. When the 100 metagenomic BGCs were aligned to their congeners in cultivated isolates, they could be collapsed into 46 unique BGCs. Thus, using two different approaches, we could accurately estimate that the 401 metagenomic BGCs of all GCFs represent ∼200 actual BGCs in the shipworm gills. To the best of our knowledge, this type of estimate has not been possible for other metagenomes/symbioses and represents a uniquely powerful aspect of this system.

Only 8 GCFs are widely distributed in 10 or more isolates, and these are mostly pathways that are universal or nearly universal in *T. turnerae*, which is overrepresented in our data set (Figs. 7 and 8). By contrast to isolate genomes in which we found many GCFs that occur in only a single genome, in the metagenomes most of the 107 GCFs are found in multiple specimens. Forty-five GCFs are found in multiple species of shipworms. Sixty-two GCFs were only found in a single shipworm species; 26 of these were only found in a single specimen (Fig. 5). These data demonstrate that accessing diverse shipworm specimens, as well as diverse shipworm species, will lead to the discovery of many novel BGCs. In addition, this result reinforces the strain-level variation found in shipworms revealed both in metagenome assembly results as well as in DNA gyrase B SNP analysis.

**Figure 7.**
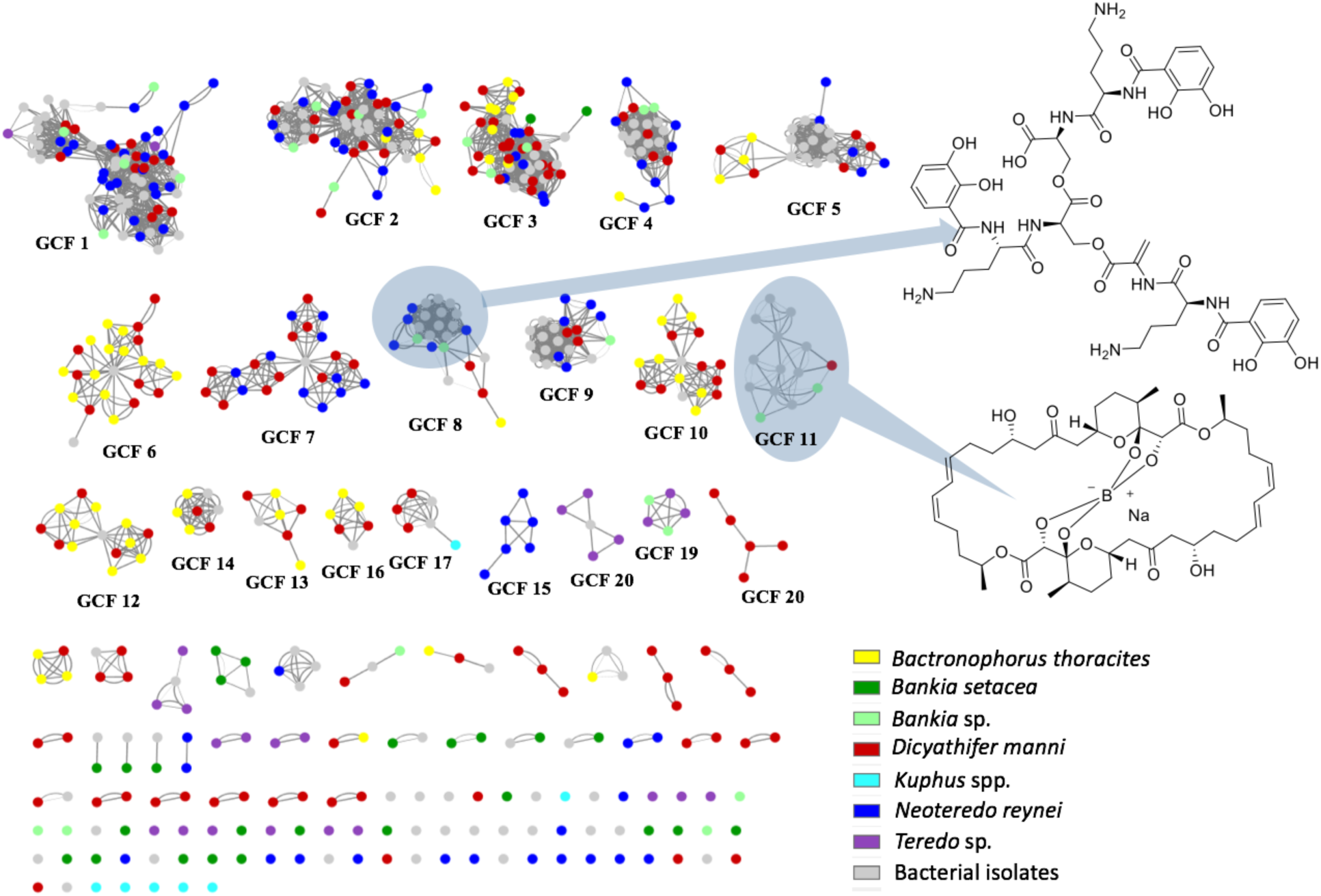
GCF distribution across shipworm species. Shown is a similarity network diagram, in which circles indicate individual BGCs from sequenced isolates (gray) and gill metagenomes (colors indicate species of origin; see legend). Lines indicate the MultiGeneBlast scores between identified BGCs, with thinner lines indicating a lower degree of similarity. For example, the cluster labeled “GCF_8” encodes the pathway for the siderophore turnerbactin, the structure of which is shown at right. The main cluster, circled by a light blue oval, includes BGCs that are very similar to the originally described turnerbactin gene cluster. More distantly related BGCs, with fewer lines connecting them to the majority nodes in GCF_8, might represent other siderophores. GCF_11 likely all represent tartrolon D/E, a boronated polyketide shown at right. For detailed alignments of BGCs, see Fig. S4.

**Figure 8.**
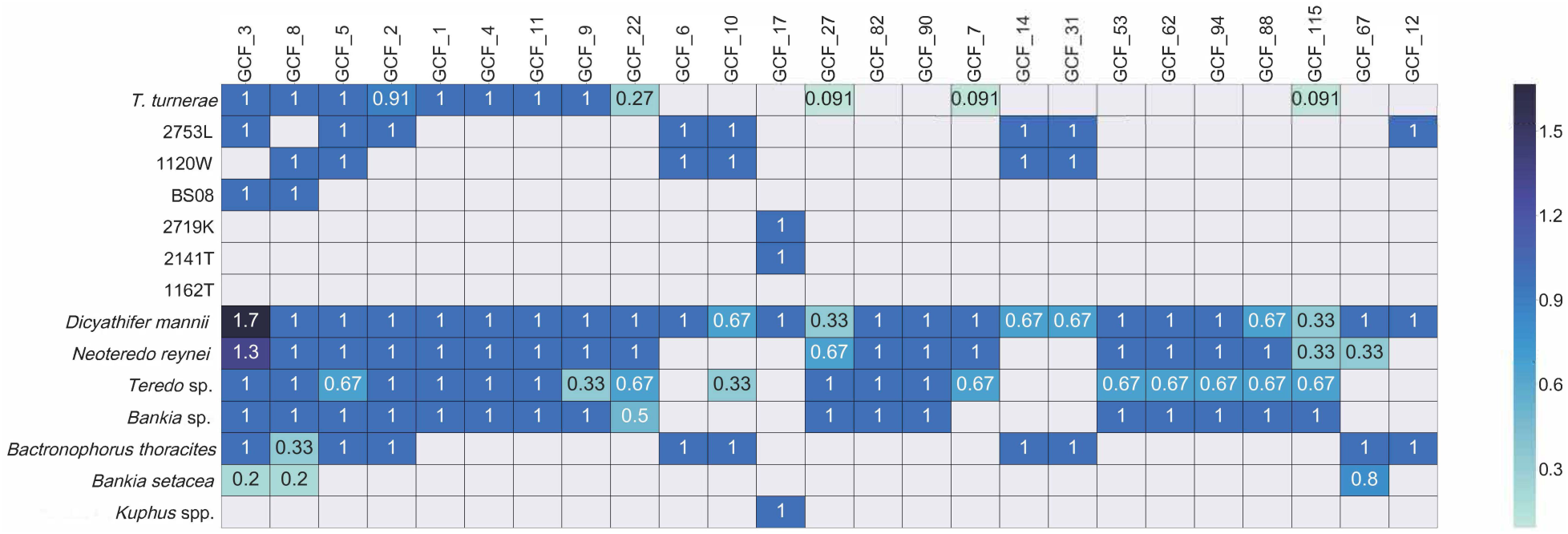
Integration of tBLASTn and networking analyses reveals the pattern of occurrence of GCFs in isolates and metagenomes. Here, we show only the most commonly occurring GCFs. The values in each box indicate the BGC occurrence per specimen for each GCF (see Fig. S5 for details). When the number equals 1, then the BGC is found in all specimens of that species. When the number is less than one, then it indicates the fraction of specimens in which the pathway is found. A number greater than one is specific to GCF_3, when two different types are possible (see Fig. 8). In that case, in two *D. mannii* specimens and one *N. reynei* specimen, there are two different classes of GCF_3, and only one class in the other specimens.

To obtain a more refined view of BGC distribution, we first used the MultiGeneBlast (31) output to construct a similarity network (Fig. 7). The network provided an easily interpretable diagram of how GCFs are distributed among bacteria. However, a notable shortcoming was observed. In a long-term drug discovery campaign, we have found the tartrolon BGC in nearly all *T. turnerae* strains ((18), unpublished observation). However, this BGC was observed in only a few of the *T. turnerae*-hosting shipworms via MultiGeneBlast. This is caused by a technical problem in assembly that we often see with large *trans*-acyltransferase (*trans*-AT) pathways from complex samples (37). Thus, we were concerned that networking might underreport the similarity of some types of biosynthetic pathways.

To remedy this problem, we obtained GCFs from cultivated isolates and searched them against metagenome contigs using tBLASTn (Fig. 8). This provided an orthogonal view of secondary metabolism in shipworms, revealing the presence of the tartrolon pathway, as well as other pathways that do not assemble well in metagenomes because of characteristics such as repetitive DNA sequences. A weakness of this second method is that it does not tell us whether two pathways are related enough to encode the production of similar compounds. Thus, these two methods provide different insights into BGCs in shipworm gills.

The close similarity of BGCs between cultivated isolates and metagenomes further reinforced the species identities determined by gANI (Table 1). Since secondary metabolism is often one of the most variable genomic features in bacteria, the sharing of multiple pathways between gills and isolates provides further evidence that the isolates are representative of the true symbionts found in gills.

We identified three categories of GCFs: (a) GCFs that are widely shared among shipworm species, (b) GCFs that were specific to select shipworm and symbiont species, and (c) GCFs that were distributed among specimens without obvious relationship to host or symbiont species identity. These pathways are described in the following sections.

**(a) Widely shared GCFs.** Four pathways (GCF_2, GCF_3, GFC_5, and GCF_8) were prevalent in all wood-eating shipworms, regardless of sample location (Figs. 7 and 8). These GCFs were encoded in the genomes of *T. turnerae,* the most widely distributed shipworm symbiont, and those of several other *Cellvibrionales* symbiont isolates from wood eating shipworms (especially the pathway-rich isolate 2753L).

The most widely occurring pathway in shipworm gill metagenomes is GCF_3. It was identified in all gill metagenomes with cellulolytic symbionts, including the metagenomes of specimen *B. setacea* BSG2. It occurs in all *T. turnerae* strains, as well as in *Cellvibrionales* strains 2753L and Bs08. It was first annotated as “region 1” in the *T. turnerae* T7901 genome and encodes an elaborate hybrid *trans*-AT PKS-NRPS pathway (14). Unlike all other GCFs identified in shipworm metagenomes and isolates, GCF_3 could be subdivided into at least three discrete categories, each of which included different gene content (Fig. 9). The first category, identified in *T. turnerae* T7901, encodes a PKS and a single NRPS, in addition to several potential modifying enzymes. In strain Bs08, instead of just a single NRPS, GCF_3 contains three NRPS genes. Presumably, Bs08 and T7901 produce products with similar or identical polyketides and amino acids, except that BS08 adds two more amino acids to the chain. *Cellvibrionales* 2753L encoded the third pathway type, which was similar to that found in T7901 except with different flanking genes that might encode modifying enzymes. Thus, T7901 and 2753L might make identical or very similar polyketide-peptide scaffolds, which are modified slightly differently after scaffold synthesis. The presence of a single GCF that encodes similar but non-identical products suggests a dynamic pathway evolution within shipworms.

**Figure 9.**
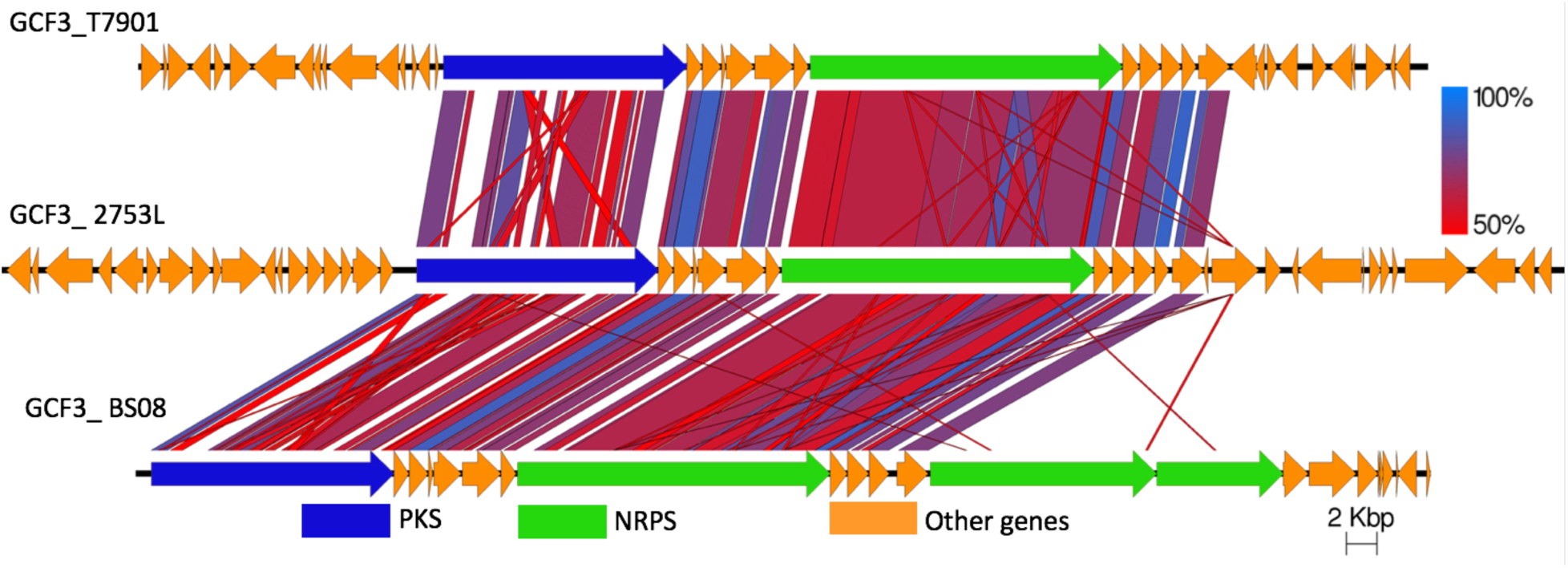
Three types of GCF_3 gene clusters are distributed in all cellulolytic shipworms in this study. tBLASTx was used to compare the clusters, demonstrating the presence of three closely related GCF_3 gene families found in all cellulolytic shipworm gills.

GCF_2 encodes a NRPS / *trans*-AT PKS pathway, the chemical products of which are unknown. It is found in all shipworm specimens in this study and in all *T. turnerae* strains. It is also present in *Cellvibrionales* strain 2753L. This explains its presence in *B. thoracites* despite the absence of *T. turnerae* in this species. GCF_2 is synonymous with “region 3” described in the annotation of the *T. turnerae* T7901 genome (14).

GCF_5 encodes a combination of terpene cyclase and predicted arylpolyene biosynthetic genes, which were unrecognized in the initial sequence analysis of *T. turnerae* T7901 (14), since the arylpolyene pathways are recent discoveries (38). Although the cyclase and surrounding regions have all of the genes necessary to make and export hopanoids, the GCF_5 biosynthetic product is unknown. In addition to occurring in all *T. turnerae* strains, GCF_5 is present in *Cellvibrionales* strains 1120W and 2753L. The pathway was detected in all wood-eating specimens except *Teredo* sp. TBF07 (Fig. 9).

GCF_8 is exemplified by the previously described turnerbactin BGC, from *T. turnerae* T7901. Turnerbactin is a catecholate siderophore, crucial to iron acquisition in *T. turnerae* (17). The BGC for turnerbactin was identified and described as “region 7” in the previously published *T. turnerae* T7901 genome. GCF_8 is present in all *T. turnerae* genomes sequenced here. Other *Cellvibrionales* strains, including 2753L from *B. thoracites* and Bs08 from *B. setacea (*neither of which contains *T. turnerae)*, also encode turnerbactin-like siderophore synthesis. GCF_8 was also found in the metagenome of one specimen of *B. thoracites*. Beyond bacterial iron acquisition, siderophores are also important in strain competition and potentially in host animal physiology (39, 40), possibly explaining the widespread distribution of GCF_8. From the clustering pattern in Fig. 7, it is likely that GCF_8 comprises at least three different, but related types of gene clusters. Thus, GCF_8 likely represents catecholate siderophores, but not necessarily turnerbactin.

**(b) Bacterial species-specific GCFs.** In addition to the four GCFs described above that have a wide distribution, GCFs 1, 4, and 11 were found in all *T. turnerae*-containing shipworms. GCF_1 is a *trans*-AT PKS-NRPS pathway that appears to be split into two clusters in some shipworm isolates, including *T. turnerae* T7901, in which it was previously annotated as “region 4” and “region 5”. GCF_4 is the previously described “region 8” PKS-NRPS from *T. turnerae* T7901.

Most notably, GCF_11 encodes tartrolon biosynthesis (18). Tartrolon is an antibiotic and potent antiparasitic agent isolated from culture broths of *T. turnerae* T7901 (18, 41, 42). It has also been identified in the cecum of the shipworm. It was proposed that the bacteria synthesize tartrolon in the gill, and it is transferred to the cecum where it may play a role in keeping the digestive tract free of bacteria (18).

The gill metagenomes of *D. mannii* and *B. thoracites* indicate the abundant presence of 2753L- like strains. Like *T. turnerae* T7901, the 2753L isolate genome encodes GCFs 2, 3, and 5. However, 2753L contains several GCFs not found in *T. turnerae*, including GCFs 6, 10, 12, 13, 14, 16, 30, and 31 (listed in order of their relative frequency of occurrence in samples). All of these GCFs are also evident in *D. mannii* and *B. thoracites* gill metagenomes. These are PKS and NRPS clusters that lack close relatives according to antiSMASH annotation and thus have a potential to synthesize novel secondary metabolite classes.

Brazilian shipworms *Bankia* sp. and *Teredo* sp. contain *T. turnerae* and the major pathways found in *T. turnerae*, but they are dominated by symbiont genomes from other symbiotic *Cellvibrionaceae* bacteria. Although those species are not represented in our current culture collection, they are closely related to isolate 1162T from a Philippine specimen of *Lyrodus* sp. The metagenomes of *Bankia* sp. and *Teredo* sp. contain many GCFs that are not found in sequenced isolates (Fig. S5B). In addition, the GCFs found in *Bankia* sp. and *Teredo* sp. are not completely overlapping, implying that the *Cellvibrionaceae* bacteria found in these different host species are distinct. A high number of GCFs were found, indicating that potentially the symbionts might have a similar GCF content as the GCF-rich isolate 2753L.

The *B. setacea* specimens shown in Fig. S5B contain pathways specifically found in *Cellvibrionaceae* isolate BSC2, which is the major bacterium observed in the *B. setacea* gill metagenome sequences.

The *K. polythalamius* gill metagenome and its cultivated sulfur oxidizing symbiont *T. teredinicola* contain relatively few BGCs, but strikingly two NRPS-containing GCFs have been found in all shipworm specimens containing the sulfide-oxidizing symbionts (*K. polythalamius* and *D. mannii*) and all sulfide-oxidizing symbiont isolates (*T. teredinicola* and isolate 2719K). One of these, GCF_17, is shown in Fig. 8. Based on our analyses, it is clear that the cellulolytic symbionts contain more abundant and diverse BGCs.

**(c) GCFs for which patterns of occurrence are not obviously related to host species identity.** Overall, the most abundant pathways in shipworms were identical to those from the cultivated isolate genomes that were mapped to each shipworm metagenome (Figs. 7 and 8). Since specific bacterial symbionts are distributed among shipworm hosts in patterns that are predicted by host species identity and life habits, the presence of abundant GCFs also follow similar patterns. However, as described above, many pathways were found only once or occurred relatively rarely among symbiont genomes and gill metagenomes. In these cases, trends of host symbiont co-occurrence could not be discerned. This trend is reinforced in Fig. 7, where most GCFs in the diagram occur only once (single, unlinked spots). Thus, while the occurrence of several biosynthetic pathways is evolutionarily conserved among host species and thus likely have a uniquely critical role in the symbiosis, most are not conserved. These observations suggest that more comprehensive sampling of shipworm specimens, species, and cultivated isolates will yield many additional, unanticipated BGCs.

### Variability in conserved shipworm GCFs increases potential compound diversity

Even among conserved GCFs, some variability was observed. This is evident in the BGC network analysis shown in (Fig. 7), where subclusters indicate slightly different GCF organization. For example, in the ubiquitous GCF_3, the three different pathway variants appear in the network as bulges within the cluster. The siderophore pathway GCF_8 contains one central cluster, encoding turnerbactin pathways, and an extended arm that appears to encode compounds that are related to, but not identical to, turnerbactin. Thus, the shape of the network clusters indicates the potential chemical diversity encoded in individual GCFs.

## Conclusions

In shipworms, cellulolytic bacteria were long known to specifically inhabit gills and were hypothesized to be the cause of an evolutionary path that leads to wood-specialization in most of the family, along with drastic morphophysiological modifications (1, 5, 43). These symbionts could be cultivated, although only recently have we been able to sample the full spectrum of major symbionts present in gills. The unexpected finding that *T. turnerae* T7901 was exceptionally rich in BGCs – proportionately denser in BGC content than *Streptomyces* spp. (14, 16) – led us to investigate shipworms as a source of new bioactive compounds.

Here, we show that cultivated isolates obtained from shipworm gills accurately represent the bacteria living within the gills. They are the same species, and often are nearly identical at the strain level. They contain many of the same BGCs. The gills of shipworms contain about 1-3 major species of symbiotic bacteria, along with a small percentage of other less consistently occurring bacteria. Complicating this relatively simple picture, there is significant strain variation within shipworms. The observed symbiont species mixtures are representative of the animal lifestyles. For example, *K. polythalamius* appears to thrive entirely on sulfide oxidation (7), as required in its sediment habitat, while the other shipworms contain various cellulolytic bacteria responsible for wood degradation. *D. mannii* likely has a more complex lifestyle, since it contains the sulfur-oxidizing bacterium strain 2719K and the cellulolytic species *T. turnerae* and strain 2753L.

The key finding is that BGCs in the metagenomes are represented in the strains in our culture collection. This is a rare event in the biosynthetic literature. In most other marine systems, it has been very challenging to cultivate the symbiotic bacteria responsible for secondary metabolite production (44). In some organisms, such as humans, there are many representative cultivated isolates that produce secondary metabolites, but connecting those metabolites to human biology, or even to their existence in humans, is quite challenging (16, 45). Here, we have defined an experimentally tractable system to investigate chemical ecology that circumvents these limitations. Our results reveal potentially important chemical interactions that would affect a variety of marine ecosystems and a novel and underexplored source of bioactive metabolites for drug discovery.

It has not escaped our notice that this work provides the foundation for understanding the connection between symbiont community composition, secondary metabolite complement, and host lifestyle and ecology. It has proven difficult to link these factors together in relevant models. The existence of aquaculture and transformation methods for shipworms and their symbiotic bacteria will enable a rigorous, hypothesis-driven understanding of the role of complex metabolism in symbiosis.

## Methods

### Collection and processing of biological material

Shipworm samples (Table S1) were collected from found wood. Briefly, infested wood was collected and transported immediately to the laboratory or stored in the shade until extraction (< 1 day). Specimens were carefully extracted to avoid damage using woodworking tools. Extracted specimens were processed immediately or stored in individual containers of filtered seawater at 4 °C until processing. Specimens were checked for viability by siphon retraction in response to stimulation and observation of heartbeat, and live specimens selected. Specimens were assigned a unique code, photographed and identified. Specimens were dissected using a dissecting stereoscope. Taxonomic vouchers (valves, pallets, and siphonal tissue for sequencing host phylogenetic markers), were retained and stored in 70% ethanol. The gill was dissected, rinsed with sterile seawater, and divided for bacterial isolation and metagenomic sequencing. Once the gill was dissected it was processed immediately or flash-frozen in liquid nitrogen.

Of the animals that we obtained in field collections we analyzed three specimens each of *Bactronophorus thoracites*, *Kuphus* spp., *Neoteredo reynei*, and *Teredo* sp., two specimens of *Bankia* sp., and five specimens of *Bankia setacea*. These animals were divided into three geographical regions (Fig. 1): the Philippines (*B. thoracites* and *D. mannii* from Infanta, Quezon; *Kuphus* spp. from Mindanao and Mabini); Brazil (*N. reynei* from Rio de Janeiro, *Teredo* sp., and *Bankia* sp. from Ceará); and the United States (*B. setacea*). The purpose of sampling this range was to determine whether there are any geographical differences in gill symbiont occurrence. Most of the shipworms were obtained from mangrove wood, with the exception of *B. setacea* from unidentified found wood, and *Kuphus* spp. from both found wood and mud.

### Bacterial isolation, DNA extraction and analysis

*Teredinibacter turnerae* strains (with T prefix) were isolated using the method described in Distel el al. 2002 (13), while *Bankia setacea* symbionts (with Bs prefix) were obtained using the technique indicated in O’Connor et al. 2014 (9). Sulfur-oxidizing symbionts were isolated using the protocol specified in Altamia et al. 2019 (22). For this study, additional *T. turnerae* and novel cellulolytic symbionts from Philippine specimens (with prefix PMS) were isolated (Table S1). Briefly, dissected gill were homogenized in sterile 75% natural seawater buffered with 20 mM HEPES, pH 8.0 using a Dounce homogenizer. Tissue homogenates were either streaked on shipworm basal medium cellulose (5) plates (1.0% Bacto Agar) or stabbed into soft agar (0.2% Bacto Agar) tubes and incubated at 25 °C until cellulolytic clearings developed. Cellulolytic bacterial colonies were subjected to several rounds of restreaking to ensure clonal selection. Contents of soft agar tubes with clearings were streaked on fresh cellulose plates to obtain single colonies. Pure colonies were then grown in 6 mL SBM cellulose liquid medium in 16 × 150 mm test tubes until the desired turbidity was observed. For long-term preservation of the isolates, a turbid medium was added to 40% glycerol at 1:1 ratio and frozen at −80 °C. Bacterial cells in the remaining liquid medium were pelleted by centrifugation at 8,000 *g* and then subjected to genomic DNA isolation. The small-subunit ribosomal (SSU) 16S rRNA gene of the isolates was then PCR amplified using 27F (5’-AGAGTTTGATCCTGGCTCAG-3’) and 1492R (5’-GGTTACCTTGTTACGACTT-3’) from the prepared genomic DNA and sequenced. Phylogenetic analyses of 16S rRNA sequences was performed using programs implemented in Geneious, version 10.2.3. Briefly, sequences were aligned using MAFFT (version 7.388) by using the E-INS-i algorithm. The aligned sequences were trimmed manually, resulting in a final aligned dataset of 1,125 nucleotide positions. Phylogenetic analysis was performed using FastTree (version 2.1.11) using the GTR substitution model with optimized Gamma20 likelihood and rate categories per site set to 20.

Genomic DNA used for whole genome sequencing of novel isolates and select *T. turnerae* strains were prepared using CTAB/phenol/chloroform DNA extraction method detailed in https://www.pacb.com/wp-content/uploads/2015/09/DNA-extraction-chlamy-CTAB-JGI.pdf.

The purity of the extracted genomic DNA was then assessed spectrophotometrically using Nanodrop and the quantity was estimated using agarose gel electrophoresis. Samples that passed the quality control steps were submitted to Joint Genome Institute – Department of Energy (JGI-DOE) for whole genome sequencing. The sequencing platform and assembly method used to generate the final isolate genome sequences used in this study are detailed in Table S1 A.

### Metagenomic DNA extraction

Gill tissue samples from Philippine shipworm specimens (Table S1 B) were flash-frozen in liquid nitrogen and stored at −80°C prior to processing. Bulk gill genomic DNA was purified by Qiagen Blood and Tissue Genomic DNA Kit using the manufacturer’s suggested protocol.

Gill tissue samples from Brazil shipworm specimens were pulverized by flash-freezing in liquid nitrogen and submitted to metagenomic DNA purification by adapting a protocol previously optimized for total DNA extraction from cnidaria tissues (46, 47). Briefly, shipworms gills were carefully dissected (taking care not to get intersections with other organs), submitted to a series of five washes with 3:1 sterile seawater / distilled water for removal of external contaminants, and macerated until powdered in liquid nitrogen. Powdered tissues (∼150 mg) were then transferred to 2 mL microtubes containing 1 mL of lysis buffer [2% (m/v) cetyltrimethyl ammonium bromide (Sigma Aldrich), 1.4 M NaCl, 20 mM EDTA, 100 mM Tris-HCl (pH 8.0), with freshly added 5 μg proteinase K (v/v; Invitrogen), and 1% 2-mercaptoethanol (Sigma Aldrich)] and submitted to five freeze-thawing cycles (−80 °C to 65 °C). Proteins were extracted by washing twice with phenol:chloroform:isoamyl alcohol (25:24:1) and once with chloroform. Metagenomic DNA was precipitated with isopropanol and 5 M ammonium acetate, washed with 70% ethanol, and eluted in TE buffer (10 mM Tris-HCl, 1 mM EDTA). Metagenomic libraries were prepared using the Nextera XT DNA Sample Preparation Kit (Illumina) and sequenced with 600-cycle (300 bp paired-end runs) MiSeq Reagent Kits v3 chemistry (Illumina) at the MiSeq Desktop Sequencer.

### Metagenome sequencing and assembly

Five *Bankia setacea* metagenome sequencing raw read files were obtained from the JGI database and reassembled using the methods described below (for accession numbers, see Table S1 A). *Kuphus polythalamius* gill metagenomes (KP2132G and KP2133G) were obtained from a previous study (7). Metagenomes from *Kuphus* sp. specimen KP3700G and *Dicyathifer mannii* and *Bactronophorus thoracites* specimens were sequenced using an Illumina HiSeq 2000 sequencer with ∼350 bp inserts and 125 bp paired-end runs at the Huntsman Cancer Institute’s High Throughput Genomics Center at the University of Utah. Illumina fastq reads were trimmed using Sickle (48) with the parameters (pe sanger -q 30–l 125). The trimmed FASTQ files were converted to FASTA files and merged using the Perl script ‘fq2fq’ in IBDA_ud package (49). Merged FASTA files were assembled using IDBA_ud with standard parameters in the Center for High Performance Computing at the University of Utah. For metagenome samples from Brazil, all *Neoterdo reynei* gill metagenomic samples previously analyzed were re-sequenced here to improve coverage depth (26). *Teredo* sp. and *Bankia* sp. gill metagenomes were sequenced using Illumina Miseq. The raw reads were assembled using either the metaspades pipeline of SPAdes (50, 51) or IDBA-UD (49). Before assembly, raw reads were merged using BBMerge (52). Non-merged reads were filtered and trimmed using FaQCs (53).

### Identification of bacterial sequences in metagenomic data

Assembly-assisted binning was used to sort and analyze trimmed reads and assembled contigs into clusters putatively representing single genomes using MetaAnnotator (54). Each binned genome was retrieved using Samtool (55, 56). To identify bacterial genomes, genes for each bin were identified with Prodigal (57). Protein sequences for bins with coding density >50% were searched against NCBI nr database with DIAMOND (58). Bins with 60% of the genes hitting bacterial subject in the nr database were considered to originate from bacteria.

For *B*. *setacea* metagenome samples and the ones from Brazil, structural and functional annotations were carried out using DFAST (59), including only contigs with length ≥500 bp. All metagenomes were binned using Autometa (60). First, each contig’s taxonomic identity was predicted using make_taxonomy_table.py, including only contigs ≥1000 bp. Predicted bacterial and archaeal contigs were binned (with recruitment via supervised machine learning) using run_autometa.py.

### gANI comparison and reads counts calculation

Each bacterial bin was compared to the 23 shipworm isolate genomes using gANI and AF values (61). With a cut-off of AF >0.5 and gANI >0.9, the bacterial bins from each metagenome were mapped to cultivated bacterial genomes, and cultivated bacterial genomes were mapped against each other (Table S2). The major but not mapped bins in each genome were classified using gtdb-tk (62). The read counts for each mapped bin were either retrieved from MetaAnnotator output or calculated using bbwrap.sh (<u>sourceforge.net/projects/bbmap/</u>) with the parameters: kfilter=22 subfilter=15 maxindel=80.

### Building BGC similarity networks

BGCs were predicted from the bacterial contigs of each metagenome and from cultivated bacterial genomes using antiSMASH 4.0 (30). From the predictions, only BGCs for PKSs, NRPSs, siderophores, terpenes, homoserine lactones, and thiopeptides (as well as combinations of these biosynthetic enzyme families) were included in succeeding analyses. An all-versus-all comparison of these BGCs was performed using MultiGeneBlast (31) following the protocol previously reported (63). Bidirectional MultiGeneBlast BGC-to-BGC hits were considered to be reliable. In metagenome data, some truncated BGCs only showed single-directional correlation to a full length BGC. Those single-directional hits were refined as follows: protein translations of all coding sequences from the BGCs were compared in an all-versus-all fashion using blastp search. Only protein hits that had at least 60% identity and at least 80% coverage to both query and subject were considered as valid hits. A single-directional MultiGeneBlast BGC-to-BGC hit was retained if there were at least *n*-2 number of proteins (*n* is the number of proteins in the truncated BGC) passing the blastp refining. The remaining MultiGeneBlast hits were used to construct a network in Cytoscape (64). Finally, each BGC cluster (GCF) that had relative low number of bidirectional correlations were manually checked by examining the MultiGeneBlast alignment.

### Occurrence of GCFs in metagenomes

Based on the GCFs identified in previous step, the core biosynthetic proteins from each GCF were extracted and queried (NCBI tblastn) against each metagenome assembly. A threshold of query coverage of >50% and identity > 90% was applied to remove the nonspecific hits, and the remaining hits, in combination with the MultiGeneBlast hits, were used to make the matrix of GCFs occurrence in metagenomes.

## Acknowledgments

All collections followed Nagoya Protocol requirements; Brazilian sampling were performed under SISBIO license number 48388, and genetic resources accessed under the authorization of the Brazilian National System for the Management of Genetic Heritage and Associated Traditional Knowledge (SisGen permit number A2F0DA0). We thank the Genomics and Bioinformatics Center of Drug Research and Development Center of Federal University of Ceara for technical support.

The work was completed under supervision of the Department of Agriculture-Bureau of Fisheries and Aquatic Resources, Philippines (DA-BFAR) in compliance with all required legal instruments and regulatory issuances covering the conduct of the research. All Philippine specimens were collected under Gratuitous Permit numbers FBP-0036-10, GP-0054-11, GP-0064-12, GP-0107-15, and GP-0140-17. We thank the governments and municipalities of the Philippines and Brazil for access and help.

This work was also supported by the National Council of Technological and Scientific Development (CNPq) (http://cnpq.br) and by the Coordination for the Improvement of Higher Education Personnel (CAPES) (http://www.capes.gov.br) under the grant numbers 473030/2013-6 and 400764/2014-8 to AETS Research reported in this publication was supported by the Fogarty International Center of the National Institutes of Health under Award Number U19TW008163. The content is solely the responsibility of the authors and does not necessarily represent the official views of the National Institutes of Health. The work was supported in part by US NOAA OER award #NA190AR0110303

## Supporting Information

**Figure S1.**
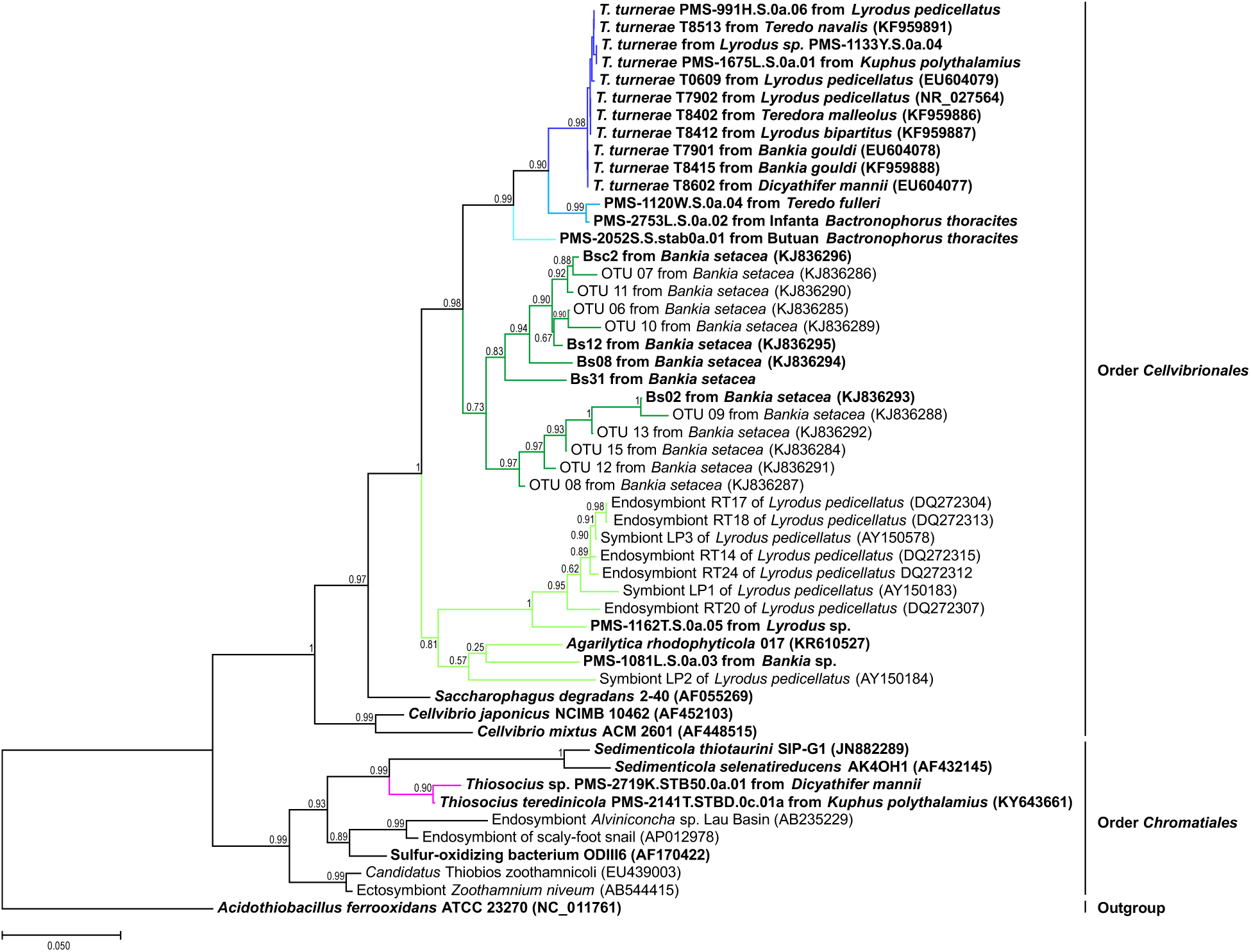
Phylogeny of shipworm gill symbionts and related free-living bacteria based on approximate maximum-likelihood tree of 16S rRNA sequences. The tree was reconstructed using 1,125 nucleotide positions employing GTR substitution model in FastTree version 2.1.11 with optimized Gamma20 likelihood and rate categories per site set to 20. Support values are indicated for each node. The scale bar represents nucleotide substitution rate per site. Cultivated shipworm symbionts and related bacteria are in boldface. The excerpted version of this tree is shown in Fig. 2A.

**Figure S2.**
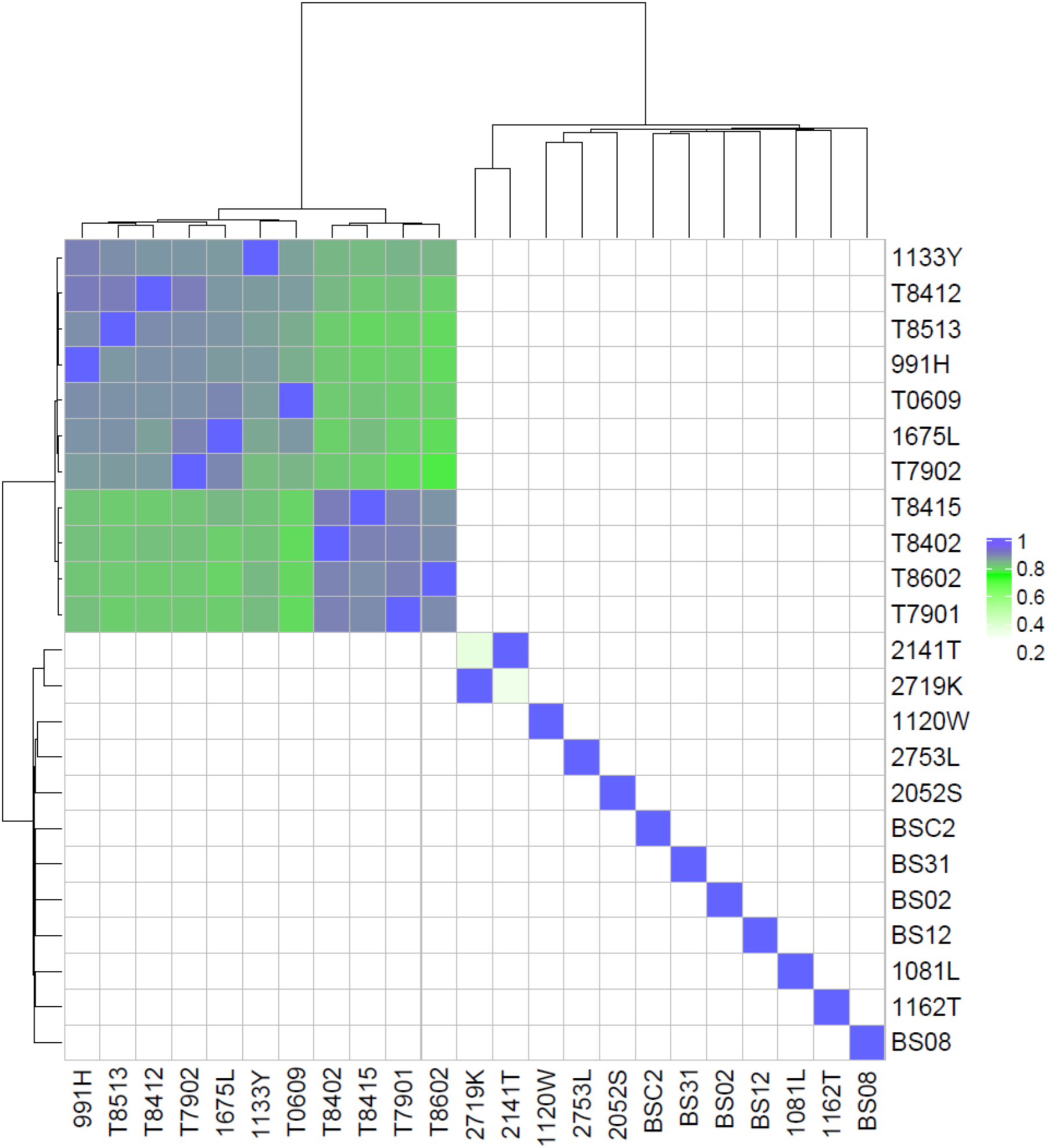
AF x gANI comparison reveals species-level differences. A heatmap with AF x gANI values comparing strain isolate genomes to each other. This analysis shows that *T. turnerae* forms 2 distinct groups, which may possibly represent different species. However, the other isolates are much more distantly related, with AF x gANI scores usually <0.2. Sulfide oxidizing bacteria also bear some similarity.

**Figure S3.**
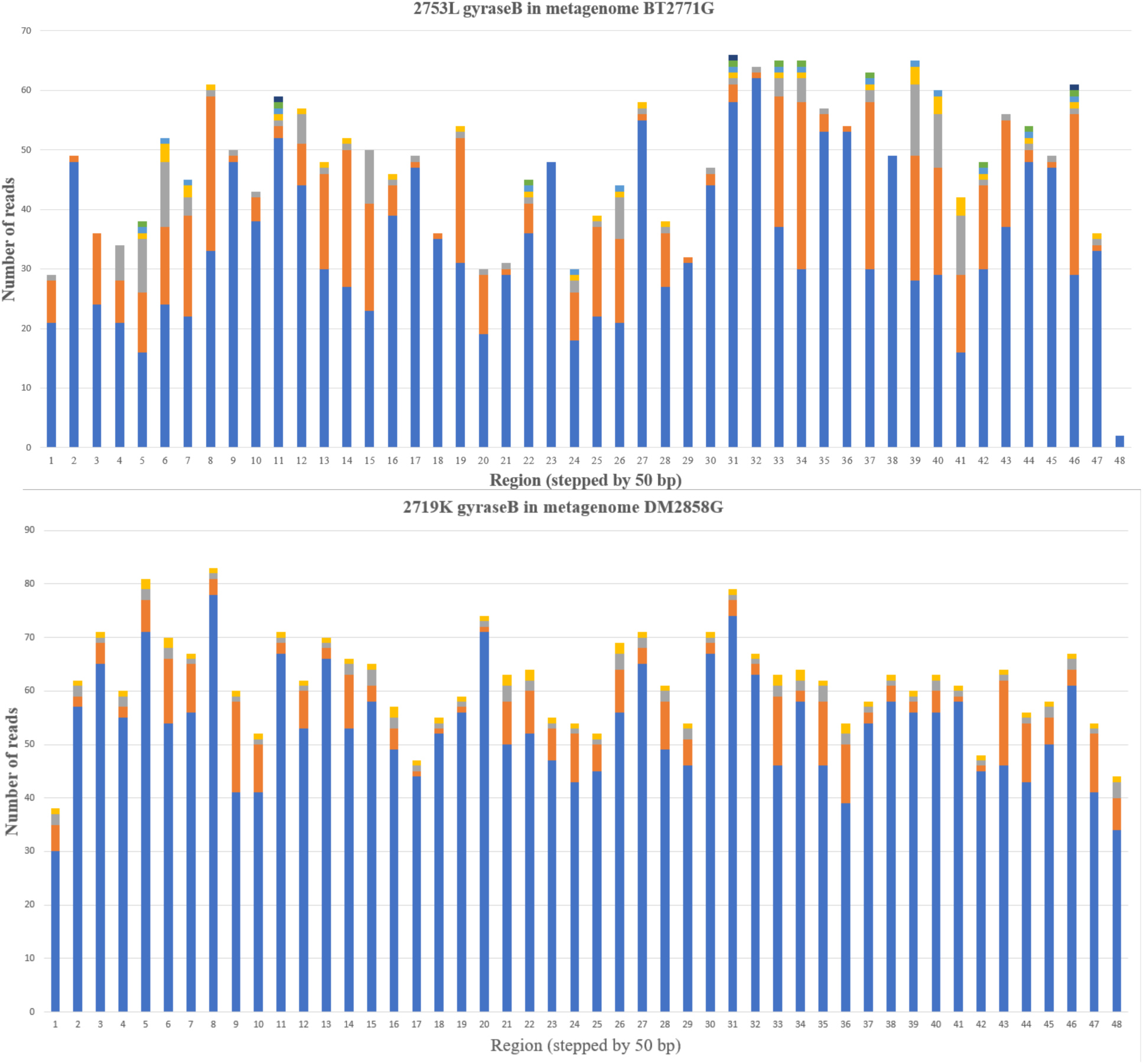
Strain variation in shipworm gill symbiont bacterial species. This figure was made as previously reported for *Kuphus* symbionts (7), using DNA gyrase B in 50 bp frames and examining SNP variation. Different colors indicate reads with different SNPs along the gyrase sequence. The *y*-axis represents number of reads observed, while the *x*-axis indicates each 50 bp region.

**Figure S4.**
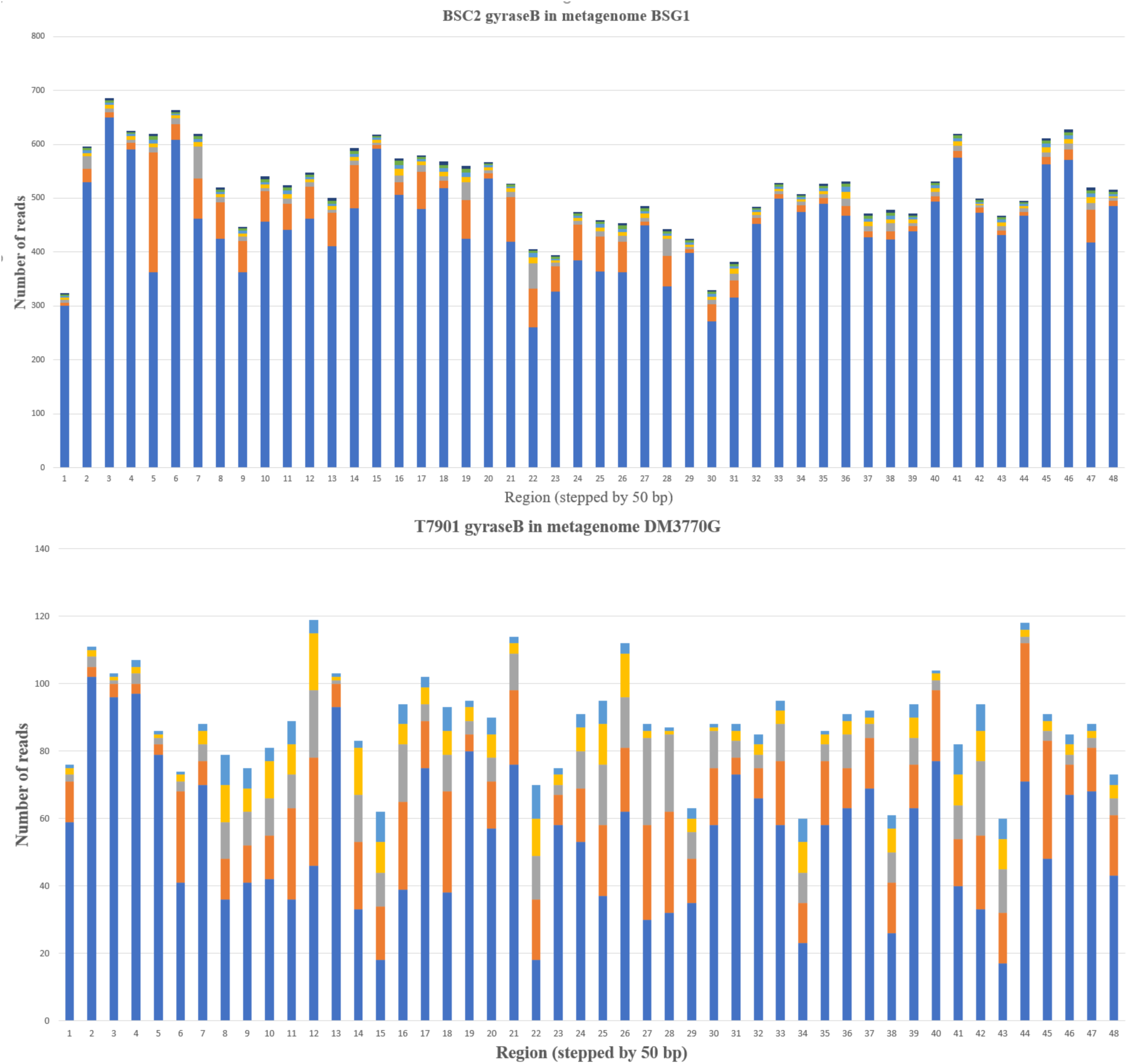
Representative alignments showing actual data underlying the clusters shown in Figs. 4, 5, 7, and 8. **A**) representative alignment of GCF_3 from genomes and metagenomes. Three subtypes were indicated by red blue and green colors; for example, the NR03 metagenome contains two copies of blue subtype. DM2858G and DM2722G contain blue and red subtypes. **B**) alignment of GCF_2. **C**) alignment of GCF_5. **D**) alignment of GCF_8.

**Figure S5.**
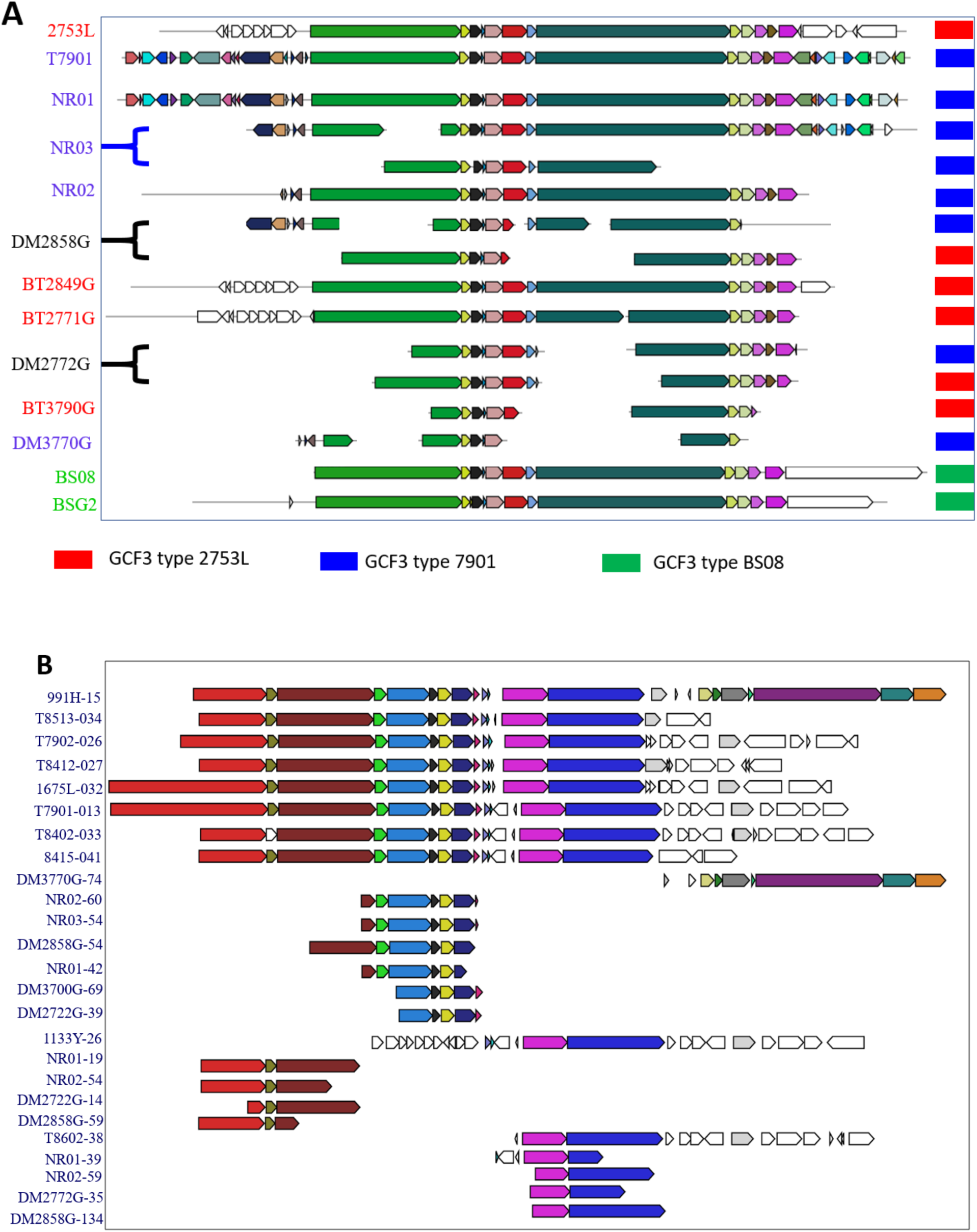
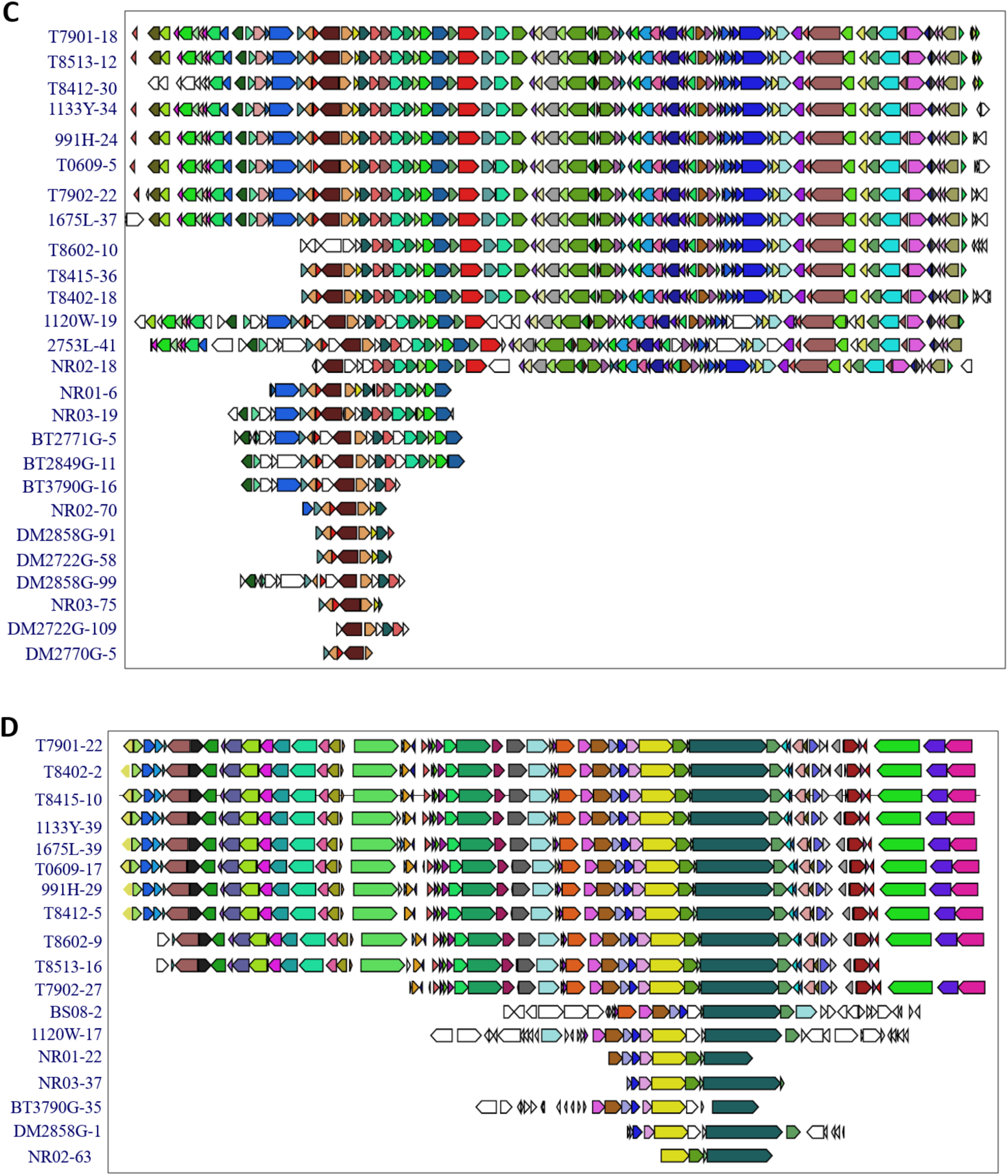
Occurrence of GCFs in individual samples, expanding what is shown in Fig. 8. **A)** GCFs found in bacterial strains. **B)** GCFs from individual shipworm specimens.

**Figure S6.**
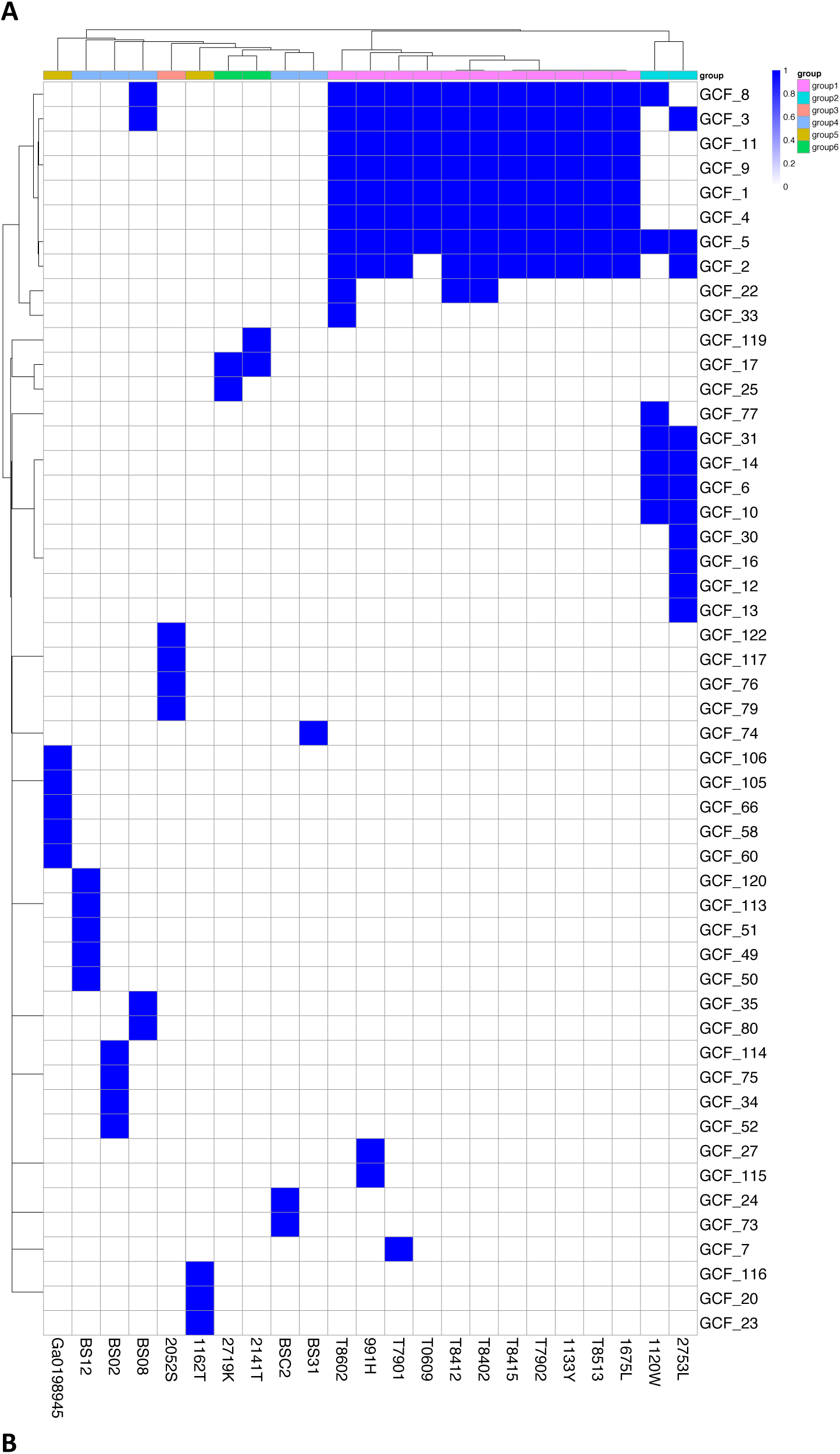
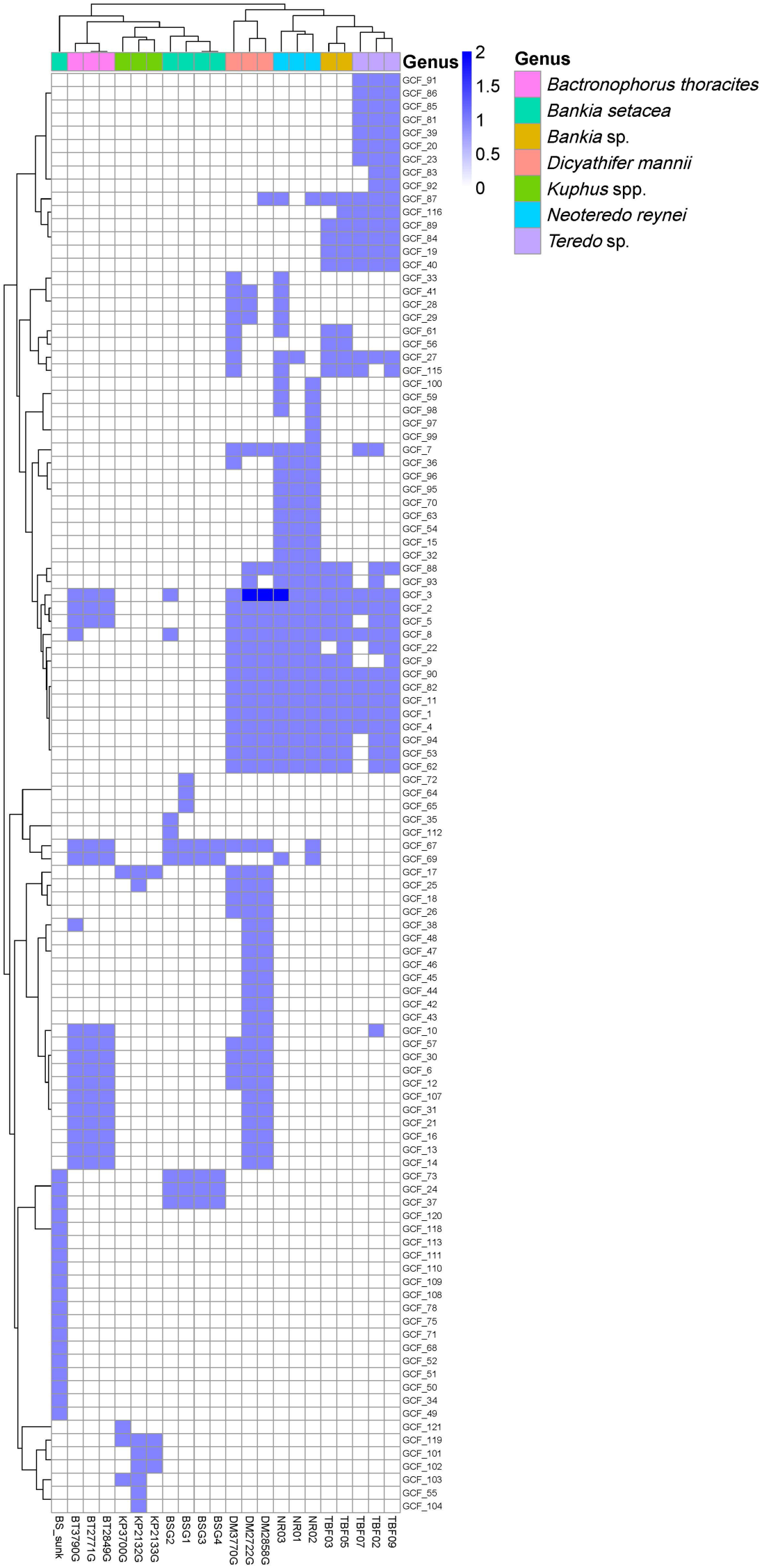
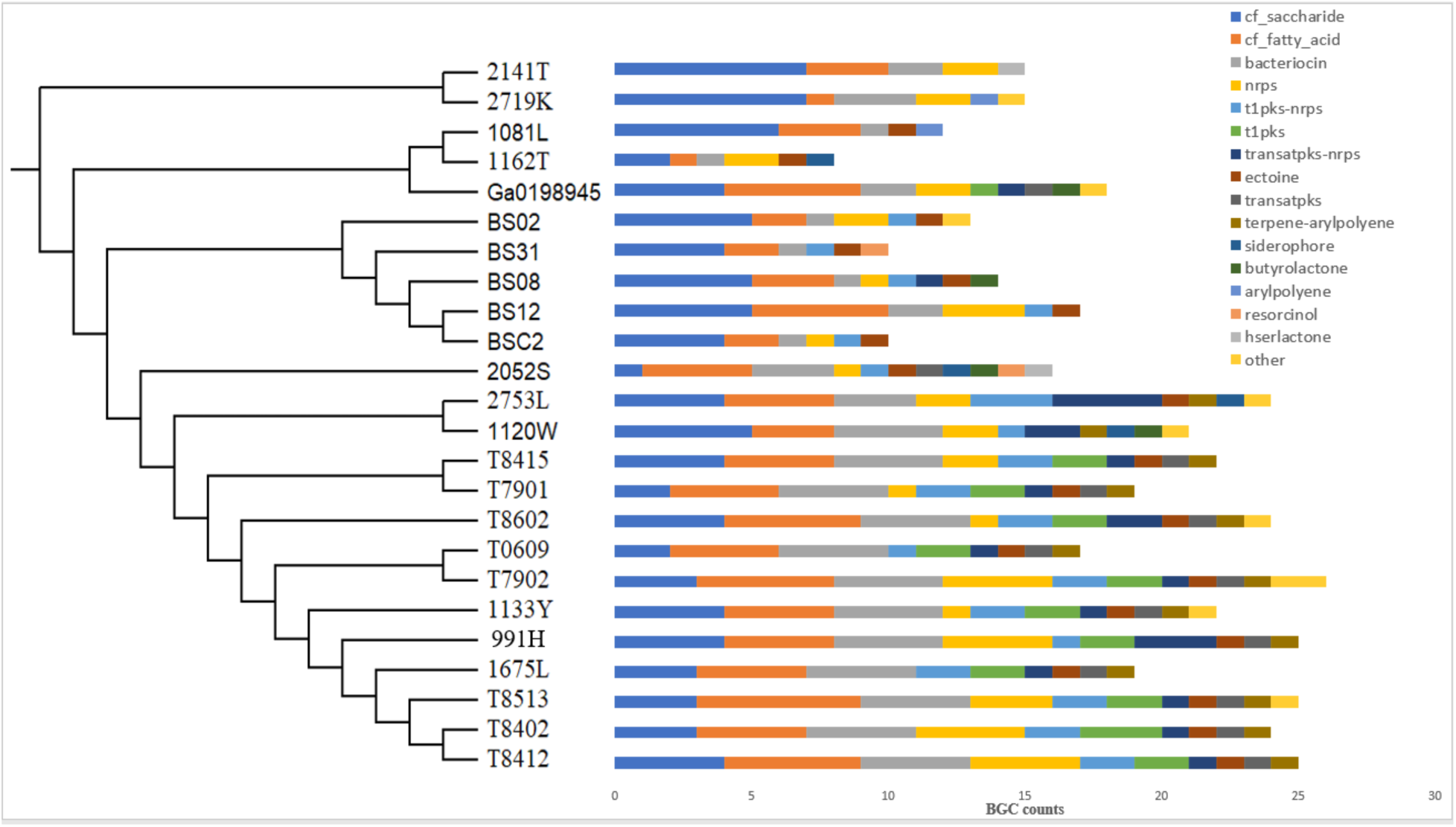
Raw antiSMASH output showing total BGCs in shipworm isolates.

**Table S1A:** Shipworm gill metagenomes used in this study.

A. Shipworm gill metagenomes used in this study.
| # | Gill metagenome | PMS-ICBG sample codes | Source shipworm species | Location | Coordinates | Sequencing center | Sequencing platform | Assembler | Reads, post trim | Size in bp | No. of contigs | N50 | %GC | IMG Genome ID | SRA accession # |
| --- | --- | --- | --- | --- | --- | --- | --- | --- | --- | --- | --- | --- | --- | --- | --- |
| 1 | DM2722G | PMS-2722P | *Dicathifer mannii* specimen PMS-2717Y | Infanta, Quezon, Philippines | N 14.68367°, E 121.63690° | Huntsman Cancer Institute, University of Utah | Illumina HiSeq 2000 | IDBA_ud | 187291588 | 1235295176 | 924064 | 2095 | 34.9 |  | SRX7665675 |
| 2 | BT2771G | PMS-2771X | *Bactronophorus thoracites* specimen PMS-2769U | Infanta, Quezon, Philippines | N 14.68367°, E 121.63690° | Huntsman Cancer Institute, University of Utah | Illumina HiSeq 2000 | IDBA_ud | 177392546 | 1056707310 | 813604 | 2023 | 35.4 |  | SRX7665685 |
| 3 | BT2849G | PMS-2849Y | *Bactronophorus thoracites* specimen PMS-2839H | Infanta, Quezon, Philippines | N 14.68367°, E 121.63690° | Huntsman Cancer Institute, University of Utah | Illumina HiSeq 2000 | IDBA_ud | 193099534 | 1059162705 | 814617 | 2024 | 35.4 |  | SRX7665686 |
| 4 | DM2858G | PMS-2858W | *Dicathifer mannii* specimen PMS-2823T | Infanta, Quezon, Philippines | N 14.68367°, E 121.63690° | Huntsman Cancer Institute, University of Utah | Illumina HiSeq 2000 | IDBA_ud | 186697500 | 1236681788 | 928980 | 2083 | 34.9 |  | SRX7665676 |
| 5 | DM3770G | PMS-3770U | *Dicathifer mannii* specimen PMS-3768S | Infanta, Quezon, Philippines | N 14.68367°, E 121.63690° | Huntsman Cancer Institute, University of Utah | Illumina HiSeq 2000 | IDBA_ud | 297553066 | 1328488478 | 1067922 | 1946 | 34.9 |  | SRX7665684 |
| 6 | BT3790G | PMS-3790S | *Bactronophorus thoracites* specimen PMS-3779S | Infanta, Quezon, Philippines | N 14.68367°, E 121.63690° | Huntsman Cancer Institute, University of Utah | Illumina HiSeq 2000 | IDBA_ud | 309554332 | 1127330840 | 927279 | 1873 | 35.4 |  | SRX7665687 |
| 10 | KP3700G | PMS-3700M | *Kuphus polythalamius* specimen PMS-3696Y (wood-boring) | Mabini, Batangas, Philippines | N 13.75843°, E 120.92586° | Huntsman Cancer Institute, University of Utah | Illumina HiSeq 2000 | IDBA_ud | 82015762 | 734092095 | 358482 | 4300 | 37.6 |  | SRX7665688 |
| 11 | KP2132G | PMS-2246K and PMS-2249P | *Kuphus polythalamius* specimen PMS-2132W (mud-dwelling) | Kalamansig, Sultan Kudarat, Philippines | N 6.53631°, E 124.048365° | Huntsman Cancer Institute, University of Utah | Illumina HiSeq 2000 | IDBA_ud | 318294870 | 772720664 | 424816 | 4530 | 37.6 |  | SRX7665689 |
| 12 | KP2133G | PMS-2157H, PMS-2116M, and PMS-2110W | *Kuphus polythalamius* specimen PMS-2133X (mud-dwelling) | Kalamansig, Sultan Kudarat, Philippines | N 6.53631°, E 124.048365° | Huntsman Cancer Institute, University of Utah | Illumina HiSeq 2000 | IDBA_ud | 329174268 | 795400237 | 500141 | 3879 | 37.4 |  | SRX7665690 |
| 13 | BSG1 | - | *Bankia setacea* | Puget Sound, Washington, USA | N 47.85072°, W 122.33843° | Joint Genome Institute - Department of Energy | Illumina HiSeq 2000 | SOAPdenovo, Newbler, and Minimus2 | 154360930 | 563042012 | 761912 | 985 | 35.0 | 3300000111 | - |
| 14 | BSG3 | - | *Bankia setacea* | Puget Sound, Washington, USA | N 47.957498°, W 122.529373° | Joint Genome Institute - Department of Energy | Illumina HiSeq 2000 | SOAPdenovo, Newbler, and Minimus2 | 144540774 | 620222960 | 648493 | 1550 | 34.9 | 3300000024 | - |
| 15 | BSG2 | - | *Bankia setacea* | Puget Sound, Washington, USA | N 47.957498°, W 122.529373° | Joint Genome Institute - Department of Energy | Illumina HiSeq 2000 | SOAPdenovo, Newbler, and Minimus2 | 180149584 | 540217764 | 793976 | 860 | 34.8 | 3300000110 | - |
| 17 | BSG4 | - | *Bankia setacea* | Puget Sound, Washington, USA | N 47.85072°, W 122.33843° | Joint Genome Institute - Department of Energy | Illumina HiSeq 2000 | SOAPdenovo, Newbler, and Minimus2 | 159707004 | 574332630 | 692986 | 1194 | 34.6 | 3300000107 | - |
| 19 | BS_sunk | - | *Bankia setacea* | Puget Sound, Washington, USA | N 47.85072°, W 122.33843° | Joint Genome Institute - Department of Energy | Illumina, 454 GS FLX Titanium | Newbler and Velvet | - | 26539887 | 38227 | 1943 | 45.2 | 2070309010 | - |
| 20 | NR01 | - | *Neoteredo reynei* | Coroa grande Mangrove - Sepetiba bay, Rio de Janeiro State, BR | 22.9081670° S 43.8756390° W | CEGENBIO | Illumina MiSeq | SPAdes | 9224156 | 313630826 | 413893 | 779 | 37.3 | - | SRX7665691 |
| 21 | NR02 | - | *Neoteredo reynei* | Coroa grande Mangrove - Sepetiba bay, Rio de Janeiro State, BR | 22.9081670° S 43.8756390° W | CEGENBIO | Illumina MiSeq | SPAdes | 18338062 | 416566737 | 468503 | 986 | 37.2 | - | SRX7665677 |
| 22 | NR03 | - | *Neoterodo reynei* | Coroa grande Mangrove - Sepetiba bay, Rio de Janeiro State, BR | 22.9081670° S<br>43.8756390° W | CEGENBIO | Illumina MiSeq | SPAdes | 13078802 | 309408486 | 414159 | 769 | 38.2 | - | SRX7665678 |
| 23 | TBF02 | - | *Teredo* sp. | Environmental Preservation Area of Pacoti river, Ceará State, Brazil | S 3.843111, W 38.422695<br>(3°50'35.2"S 38°25'21.7"W) | CEGENBIO | Illumina MiSeq | SPAdes | 3565711 | 74108472 | 236018 | 1037 | 40.1 |  | SRX7665679 |
| 24 | TBF03 | - | *Bankia* sp. | Environmental Preservation Area of Pacoti river, Ceará State, Brazil | S 3.843111, W 38.422695<br>(3°50'35.2"S 38°25'21.7"W) | CEGENBIO | Illumina MiSeq | SPAdes | 2205607 | 33524312 | 75538 | 1123 | 41.2 |  | SRX7665680 |
| 25 | TBF05 | - | *Bankia* sp. | Environmental Preservation Area of Pacoti river, Ceará State, Brazil | S 3.843111, W 38.422695<br>(3°50'35.2"S 38°25'21.7"W) | CEGENBIO | Illumina MiSeq | SPAdes | 3632367 | 107179837 | 230494 | 995 | 37.4 |  | SRX7665681 |
| 26 | TBF07 | - | *Teredo* sp. | Environmental Preservation Area of Pacoti river, Ceará State, Brazil | S 3.843111, W 38.422695<br>(3°50'35.2"S 38°25'21.7"W) | CEGENBIO | Illumina MiSeq | SPAdes | 3731031 | 78684542 | 258368 | 965 | 38.6 |  | SRX7665682 |
| 27 | TBF09 | - | *Teredo* sp. | Environmental Preservation Area of Pacoti river, Ceará State, Brazil | S 3.843111, W 38.422695<br>(3°50'35.2"S 38°25'21.7"W) | CEGENBIO | Illumina MiSeq | SPAdes | 4029653 | 108441874 | 340072 | 948 | 38.1 |  | SRX7665683 |

**Table S1B.**
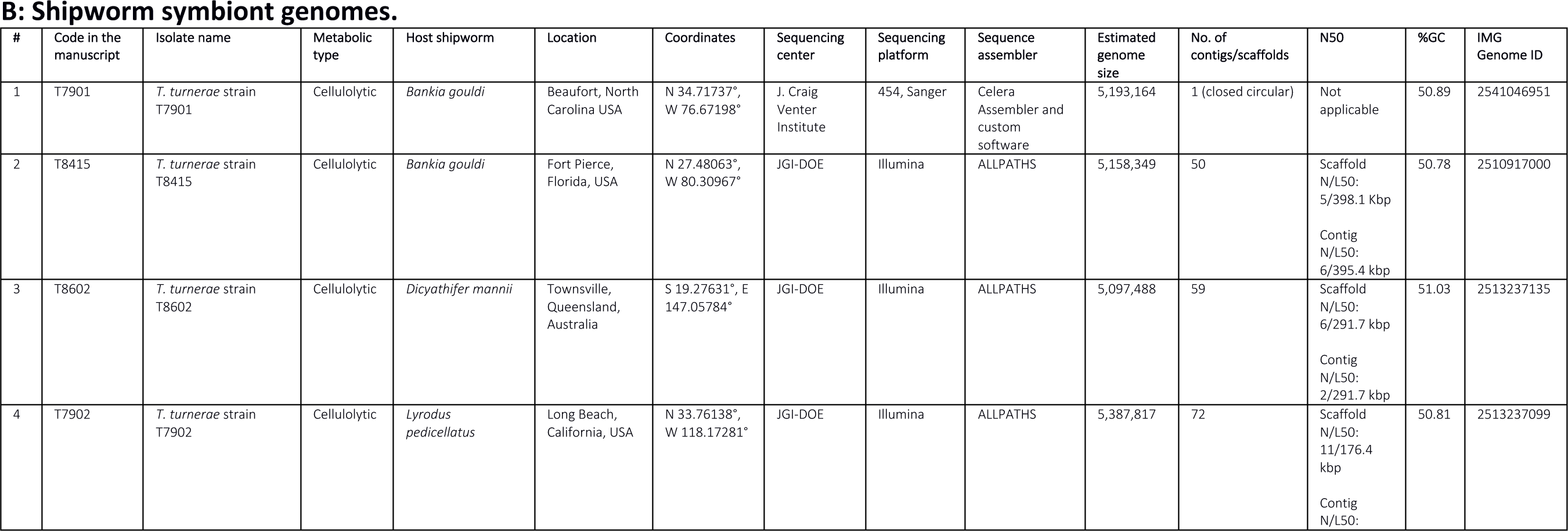
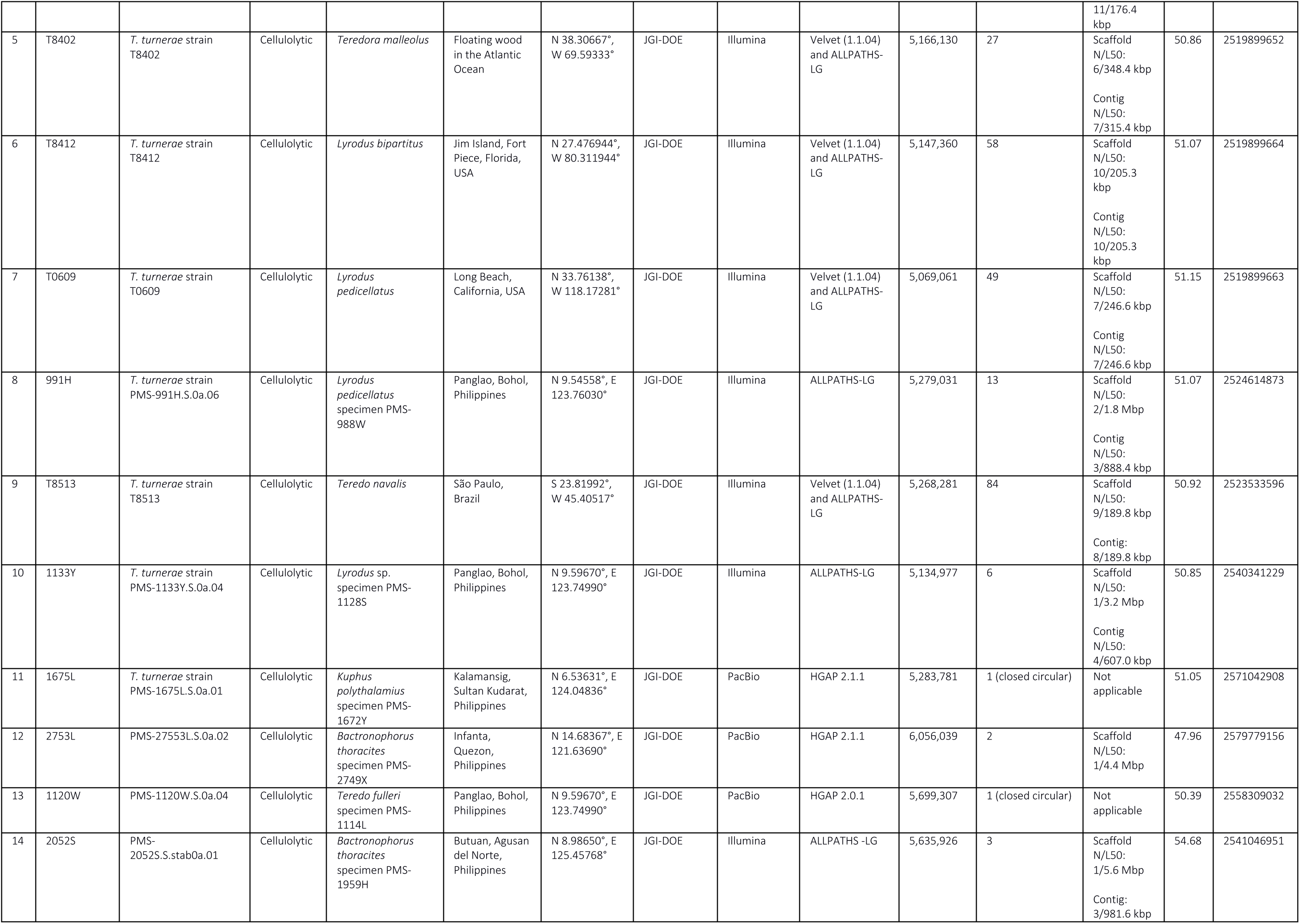
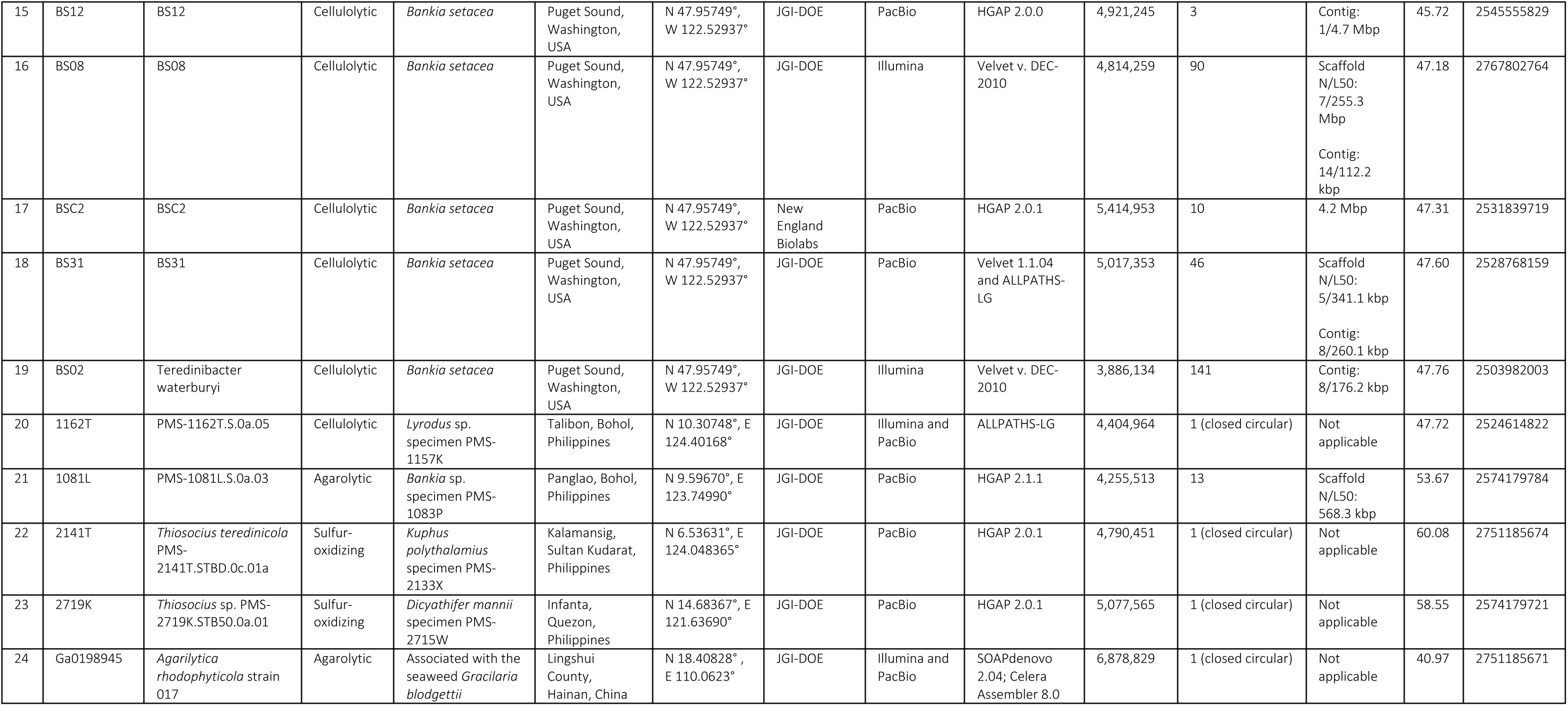
Shipworm symbiont genomes.

**Table S2:** gANI comparison of genomes and metagenome bins.

| genome_ID1 | genome_ID2 | AF | gANI | edge_score=AF*gANI | genome_length |
| --- | --- | --- | --- | --- | --- |
| 1133Y | 1675L | 0.88837 | 0.97601 | 0.867058 | 4416264 |
| 1133Y | 991H | 0.917913 | 0.976538 | 0.896377 | 4416264 |
| 1133Y | NR01_83_0.fasta | 0.625963 | 0.931249 | 0.582927 | 4416264 |
| 1133Y | NR02_1_1.fasta | 0.828551 | 0.924503 | 0.765998 | 4416264 |
| 1133Y | NR03_uc.fasta | 0.627183 | 0.936548 | 0.587387 | 4416264 |
| 1133Y | T0609 | 0.880447 | 0.974621 | 0.858102 | 4416264 |
| 1133Y | T7901 | 0.901882 | 0.930208 | 0.838938 | 4416264 |
| 1133Y | T7902 | 0.892839 | 0.976096 | 0.871497 | 4416264 |
| 1133Y | T8402 | 0.89629 | 0.931583 | 0.834969 | 4416264 |
| 1133Y | T8412 | 0.896951 | 0.972772 | 0.872529 | 4416264 |
| 1133Y | T8415 | 0.890276 | 0.932962 | 0.830594 | 4416264 |
| 1133Y | T8513 | 0.903741 | 0.975097 | 0.881235 | 4416264 |
| 1133Y | T8602 | 0.897988 | 0.930602 | 0.835669 | 4416264 |
| 1133Y | TBF05_2_0.fasta | 0.818783 | 0.968715 | 0.793167 | 4416264 |
| 1675L | 1133Y | 0.872475 | 0.975545 | 0.851139 | 4536378 |
| 1675L | 991H | 0.890956 | 0.98154 | 0.874509 | 4536378 |
| 1675L | NR01_83_0.fasta | 0.633585 | 0.931447 | 0.590151 | 4536378 |
| 1675L | NR02_1_1.fasta | 0.827928 | 0.919958 | 0.761659 | 4536378 |
| 1675L | NR03_uc.fasta | 0.635214 | 0.935692 | 0.594365 | 4536378 |
| 1675L | T0609 | 0.887587 | 0.979341 | 0.86925 | 4536378 |
| 1675L | T7901 | 0.868763 | 0.923789 | 0.802554 | 4536378 |
| 1675L | T7902 | 0.905002 | 0.986385 | 0.89268 | 4536378 |
| 1675L | T8402 | 0.86934 | 0.925605 | 0.804665 | 4536378 |
| 1675L | T8412 | 0.877281 | 0.979048 | 0.8589 | 4536378 |
| 1675L | T8415 | 0.894459 | 0.924962 | 0.827341 | 4536378 |
| 1675L | T8513 | 0.892232 | 0.982003 | 0.876175 | 4536378 |
| 1675L | T8602 | 0.855345 | 0.924807 | 0.791029 | 4536378 |
| 1675L | TBF05_2_0.fasta | 0.783679 | 0.97568 | 0.76462 | 4536378 |
| 2141T | KP2132G_543 | 0.667543 | 0.904162 | 0.603567 | 4217910 |
| 2719K | DM2722G_579 | 0.871057 | 0.994387 | 0.866168 | 4518363 |
| 2719K | DM2858G_1458 | 0.855756 | 0.991247 | 0.848266 | 4518363 |
| 2719K | DM3770G_2725 | 0.783067 | 0.976768 | 0.764875 | 4518363 |
| 2753L | BT2849G_1909 | 0.626956 | 0.988297 | 0.619619 | 5329059 |
| 2753L | DM2858G_3735 | 0.520819 | 0.985097 | 0.513057 | 5329059 |
| 991H | 1133Y | 0.888098 | 0.975928 | 0.86672 | 4574307 |
| 991H | 1675L | 0.88192 | 0.982037 | 0.866078 | 4574307 |
| 991H | NR01_83_0.fasta | 0.622196 | 0.930411 | 0.578898 | 4574307 |
| 991H | NR02_1_1.fasta | 0.8166 | 0.917586 | 0.749301 | 4574307 |
| 991H | NR03_uc.fasta | 0.632584 | 0.933684 | 0.590634 | 4574307 |
| 991H | T0609 | 0.86053 | 0.983114 | 0.845999 | 4574307 |
| 991H | T7901 | 0.873763 | 0.923005 | 0.806488 | 4574307 |
| 991H | T7902 | 0.894225 | 0.981671 | 0.877835 | 4574307 |
| 991H | T8402 | 0.880146 | 0.923846 | 0.813119 | 4574307 |
| 991H | T8412 | 0.89513 | 0.979792 | 0.877041 | 4574307 |
| 991H | T8415 | 0.869846 | 0.923357 | 0.803178 | 4574307 |
| 991H | T8513 | 0.885041 | 0.983463 | 0.870405 | 4574307 |
| 991H | T8602 | 0.85937 | 0.922279 | 0.792579 | 4574307 |
| 991H | TBF05_2_0.fasta | 0.792542 | 0.976231 | 0.773704 | 4574307 |
| BS08 | BSG2_2_0.fasta | 0.761704 | 0.992799 | 0.756219 | 4165124 |
| BSC2.fasta | BSG1_1_1.fasta | 0.87084 | 0.965521 | 0.840814 | 4577055 |
| BSC2.fasta | BSG3_2_0.fasta | 0.724032 | 0.958843 | 0.694233 | 4577055 |
| BSC2.fasta | BSG4_1_0.fasta | 0.711961 | 0.970806 | 0.691176 | 4577055 |
| BSG1_1_1.fasta | BSC2.fasta | 0.645775 | 0.973085 | 0.628394 | 6666104 |
| BSG2_2_0.fasta | BS08 | 0.920644 | 0.995137 | 0.916167 | 3513639 |
| BSG2_2_1.fasta | BSC2.fasta | 0.898617 | 0.9748 | 0.875972 | 2557003 |
| BSG2_2_4.fasta | BSC2.fasta | 0.945788 | 0.976468 | 0.923532 | 323727 |
| BSG2_2_9.fasta | BSC2.fasta | 0.904797 | 0.980741 | 0.887372 | 137905 |
| BSG3_2_0.fasta | BSC2.fasta | 0.593708 | 0.973826 | 0.578168 | 6133323 |
| BSG4_1_0.fasta | BSC2.fasta | 0.861395 | 0.974795 | 0.839684 | 3869091 |
| BT2771G_1251 | 2753L | 0.737287 | 0.985666 | 0.726719 | 931086 |
| BT2771G_1266 | 2753L | 0.959437 | 0.99215 | 0.951905 | 1872123 |
| BT2771G_2629 | 2753L | 0.903905 | 0.991576 | 0.896291 | 2667810 |
| BT2849G_1158 | 2753L | 0.784775 | 0.9885 | 0.77575 | 1191456 |
| BT2849G_1418 | 2753L | 0.990501 | 0.992228 | 0.982803 | 16107 |
| BT2849G_1523 | 2753L | 0.540321 | 0.985773 | 0.532634 | 21465 |
| BT2849G_1577 | 2753L | 0.711287 | 0.963563 | 0.68537 | 36732 |
| BT2849G_1909 | 2753L | 0.971633 | 0.991404 | 0.963281 | 3503796 |
| BT2849G_2869 | 2753L | 0.830116 | 0.988927 | 0.820924 | 687471 |
| BT3790G_1208 | 2753L | 0.745048 | 0.982692 | 0.732153 | 402075 |
| BT3790G_1493 | 2753L | 0.768579 | 0.989095 | 0.760198 | 2051745 |
| BT3790G_1981 | 2753L | 0.552787 | 0.959462 | 0.530378 | 64305 |
| BT3790G_2237 | 2753L | 0.933243 | 0.991594 | 0.925398 | 3015375 |
| BT3790G_3135 | 2753L | 0.626175 | 0.981932 | 0.614861 | 732126 |
| DM2722G_1447 | 1133Y | 0.869977 | 0.935091 | 0.813508 | 2119464 |
| DM2722G_1447 | 1675L | 0.863317 | 0.928096 | 0.801241 | 2119464 |
| DM2722G_1447 | 991H | 0.865791 | 0.927151 | 0.802719 | 2119464 |
| DM2722G_1447 | T0609 | 0.859012 | 0.928516 | 0.797606 | 2119464 |
| DM2722G_1447 | T7901 | 0.891632 | 0.983404 | 0.876834 | 2119464 |
| DM2722G_1447 | T7902 | 0.872281 | 0.927808 | 0.809309 | 2119464 |
| DM2722G_1447 | T8402 | 0.891799 | 0.981492 | 0.875294 | 2119464 |
| DM2722G_1447 | T8412 | 0.863001 | 0.926899 | 0.799915 | 2119464 |
| DM2722G_1447 | T8415 | 0.885098 | 0.981571 | 0.868787 | 2119464 |
| DM2722G_1447 | T8513 | 0.864245 | 0.927157 | 0.801291 | 2119464 |
| DM2722G_1447 | T8602 | 0.895983 | 0.985196 | 0.882719 | 2119464 |
| DM2722G_1691 | 2753L | 0.860087 | 0.991096 | 0.852429 | 2544396 |
| DM2722G_1870 | 1133Y | 0.873943 | 0.927325 | 0.810429 | 2047428 |
| DM2722G_1870 | 1675L | 0.87697 | 0.921825 | 0.808413 | 2047428 |
| DM2722G_1870 | 991H | 0.877267 | 0.920066 | 0.807144 | 2047428 |
| DM2722G_1870 | T0609 | 0.878737 | 0.922672 | 0.810786 | 2047428 |
| DM2722G_1870 | T7901 | 0.914987 | 0.981546 | 0.898102 | 2047428 |
| DM2722G_1870 | T7902 | 0.871213 | 0.920104 | 0.801607 | 2047428 |
| DM2722G_1870 | T8402 | 0.900602 | 0.980756 | 0.883271 | 2047428 |
| DM2722G_1870 | T8412 | 0.867036 | 0.919955 | 0.797634 | 2047428 |
| DM2722G_1870 | T8415 | 0.895129 | 0.981002 | 0.878123 | 2047428 |
| DM2722G_1870 | T8513 | 0.861727 | 0.920596 | 0.793302 | 2047428 |
| DM2722G_1870 | T8602 | 0.905362 | 0.982463 | 0.889485 | 2047428 |
| DM2722G_3144 | 2753L | 0.682074 | 0.989466 | 0.674889 | 3191535 |
| DM2722G_497 | 2719K | 0.707764 | 0.988715 | 0.699777 | 515829 |
| DM2722G_579 | 2719K | 0.986046 | 0.99807 | 0.984143 | 4146420 |
| DM2858G_1105 | 1133Y | 0.620173 | 0.946415 | 0.586941 | 1252896 |
| DM2858G_1105 | 1675L | 0.633836 | 0.935645 | 0.593045 | 1252896 |
| DM2858G_1105 | 991H | 0.619454 | 0.931833 | 0.577228 | 1252896 |
| DM2858G_1105 | T0609 | 0.603874 | 0.934476 | 0.564306 | 1252896 |
| DM2858G_1105 | T7901 | 0.64117 | 0.976821 | 0.626308 | 1252896 |
| DM2858G_1105 | T7902 | 0.611744 | 0.933045 | 0.570785 | 1252896 |
| DM2858G_1105 | T8402 | 0.643978 | 0.973645 | 0.627006 | 1252896 |
| DM2858G_1105 | T8412 | 0.605823 | 0.931875 | 0.564551 | 1252896 |
| DM2858G_1105 | T8415 | 0.63788 | 0.974056 | 0.621331 | 1252896 |
| DM2858G_1105 | T8513 | 0.60504 | 0.93013 | 0.562766 | 1252896 |
| DM2858G_1105 | T8602 | 0.654028 | 0.976458 | 0.638631 | 1252896 |
| DM2858G_1458 | 2719K | 0.95337 | 0.996185 | 0.949733 | 4319439 |
| DM2858G_2488 | 1133Y | 0.527697 | 0.915996 | 0.483368 | 31086 |
| DM2858G_2488 | 1675L | 0.509168 | 0.922921 | 0.469922 | 31086 |
| DM2858G_2488 | 991H | 0.519494 | 0.912874 | 0.474233 | 31086 |
| DM2858G_2488 | T7901 | 0.565238 | 0.959991 | 0.542623 | 31086 |
| DM2858G_2488 | T7902 | 0.541884 | 0.911428 | 0.493888 | 31086 |
| DM2858G_2488 | T8402 | 0.539761 | 0.97622 | 0.526925 | 31086 |
| DM2858G_2488 | T8412 | 0.533681 | 0.914949 | 0.488291 | 31086 |
| DM2858G_2488 | T8415 | 0.570546 | 0.97412 | 0.55578 | 31086 |
| DM2858G_2488 | T8602 | 0.569774 | 0.971827 | 0.553722 | 31086 |
| DM2858G_2501 | 2719K | 0.701329 | 0.98137 | 0.688263 | 411372 |
| DM2858G_2907 | 1133Y | 0.630511 | 0.914008 | 0.576292 | 2150778 |
| DM2858G_2907 | 1675L | 0.623165 | 0.90994 | 0.567043 | 2150778 |
| DM2858G_2907 | 991H | 0.622968 | 0.909982 | 0.56689 | 2150778 |
| DM2858G_2907 | T0609 | 0.626758 | 0.910536 | 0.570686 | 2150778 |
| DM2858G_2907 | T7901 | 0.652576 | 0.972146 | 0.634399 | 2150778 |
| DM2858G_2907 | T7902 | 0.624774 | 0.908152 | 0.56739 | 2150778 |
| DM2858G_2907 | T8402 | 0.648725 | 0.972033 | 0.630582 | 2150778 |
| DM2858G_2907 | T8412 | 0.630313 | 0.9102 | 0.573711 | 2150778 |
| DM2858G_2907 | T8415 | 0.646015 | 0.972657 | 0.628351 | 2150778 |
| DM2858G_2907 | T8513 | 0.618733 | 0.909773 | 0.562907 | 2150778 |
| DM2858G_2907 | T8602 | 0.641596 | 0.972786 | 0.624136 | 2150778 |
| DM2858G_3045 | 1133Y | 0.919354 | 0.933049 | 0.857802 | 1967640 |
| DM2858G_3045 | 1675L | 0.914358 | 0.926672 | 0.84731 | 1967640 |
| DM2858G_3045 | 991H | 0.916906 | 0.926561 | 0.849569 | 1967640 |
| DM2858G_3045 | T0609 | 0.916133 | 0.927309 | 0.849538 | 1967640 |
| DM2858G_3045 | T7901 | 0.932526 | 0.984445 | 0.918021 | 1967640 |
| DM2858G_3045 | T7902 | 0.907632 | 0.926104 | 0.840562 | 1967640 |
| DM2858G_3045 | T8402 | 0.928931 | 0.982109 | 0.912311 | 1967640 |
| DM2858G_3045 | T8412 | 0.910497 | 0.925235 | 0.842424 | 1967640 |
| DM2858G_3045 | T8415 | 0.928217 | 0.981995 | 0.911504 | 1967640 |
| DM2858G_3045 | T8513 | 0.908572 | 0.925417 | 0.840808 | 1967640 |
| DM2858G_3045 | T8602 | 0.934802 | 0.986011 | 0.921725 | 1967640 |
| DM2858G_3735 | 2753L | 0.861203 | 0.991039 | 0.853486 | 3409581 |
| DM3770G_1109 | T8415 | 0.528894 | 0.9172 | 0.485102 | 13290 |
| DM3770G_1432 | T7901 | 0.515957 | 0.973758 | 0.502417 | 771279 |
| DM3770G_2006 | 1133Y | 0.936908 | 0.926142 | 0.86771 | 1755333 |
| DM3770G_2006 | 1675L | 0.936995 | 0.924306 | 0.86607 | 1755333 |
| DM3770G_2006 | 991H | 0.942717 | 0.923133 | 0.870253 | 1755333 |
| DM3770G_2006 | T0609 | 0.93762 | 0.923223 | 0.865632 | 1755333 |
| DM3770G_2006 | T7901 | 0.954063 | 0.985646 | 0.940368 | 1755333 |
| DM3770G_2006 | T7902 | 0.937723 | 0.922307 | 0.864868 | 1755333 |
| DM3770G_2006 | T8402 | 0.953564 | 0.983361 | 0.937698 | 1755333 |
| DM3770G_2006 | T8412 | 0.934559 | 0.92214 | 0.861794 | 1755333 |
| DM3770G_2006 | T8415 | 0.956646 | 0.983846 | 0.941192 | 1755333 |
| DM3770G_2006 | T8513 | 0.93782 | 0.922223 | 0.864879 | 1755333 |
| DM3770G_2006 | T8602 | 0.948791 | 0.987314 | 0.936755 | 1755333 |
| DM3770G_2725 | 2719K | 0.915257 | 0.985789 | 0.90225 | 4824342 |
| DM3770G_2751 | 1133Y | 0.892276 | 0.947536 | 0.845464 | 406707 |
| DM3770G_2751 | 1675L | 0.863191 | 0.938519 | 0.810121 | 406707 |
| DM3770G_2751 | 991H | 0.875938 | 0.938369 | 0.821953 | 406707 |
| DM3770G_2751 | T0609 | 0.876034 | 0.938738 | 0.822366 | 406707 |
| DM3770G_2751 | T7901 | 0.893228 | 0.974315 | 0.870285 | 406707 |
| DM3770G_2751 | T7902 | 0.849081 | 0.937778 | 0.796249 | 406707 |
| DM3770G_2751 | T8402 | 0.882879 | 0.971073 | 0.85734 | 406707 |
| DM3770G_2751 | T8412 | 0.869786 | 0.936096 | 0.814203 | 406707 |
| DM3770G_2751 | T8415 | 0.885077 | 0.968797 | 0.85746 | 406707 |
| DM3770G_2751 | T8513 | 0.85316 | 0.936312 | 0.798824 | 406707 |
| DM3770G_2751 | T8602 | 0.901799 | 0.973795 | 0.878167 | 406707 |
| DM3770G_2901 | 2753L | 0.791226 | 0.988497 | 0.782125 | 105042 |
| DM3770G_2983 | 1133Y | 0.741582 | 0.926188 | 0.686844 | 1267746 |
| DM3770G_2983 | 1675L | 0.710092 | 0.918912 | 0.652512 | 1267746 |
| DM3770G_2983 | 991H | 0.734198 | 0.918836 | 0.674608 | 1267746 |
| DM3770G_2983 | T0609 | 0.722201 | 0.918436 | 0.663295 | 1267746 |
| DM3770G_2983 | T7901 | 0.777272 | 0.981042 | 0.762536 | 1267746 |
| DM3770G_2983 | T7902 | 0.721815 | 0.918004 | 0.662629 | 1267746 |
| DM3770G_2983 | T8402 | 0.777461 | 0.980031 | 0.761936 | 1267746 |
| DM3770G_2983 | T8412 | 0.735069 | 0.919615 | 0.67598 | 1267746 |
| DM3770G_2983 | T8415 | 0.74838 | 0.98179 | 0.734752 | 1267746 |
| DM3770G_2983 | T8513 | 0.723786 | 0.918056 | 0.664476 | 1267746 |
| DM3770G_2983 | T8602 | 0.761282 | 0.980425 | 0.74638 | 1267746 |
| DM3770G_3242 | 2753L | 0.770013 | 0.980837 | 0.755257 | 37476 |
| DM3770G_3460 | T8602 | 0.521027 | 0.958985 | 0.499657 | 36453 |
| DM3770G_5 | 2753L | 0.51835 | 0.986173 | 0.511183 | 276537 |
| DM3770G_615 | 1133Y | 0.870698 | 0.943556 | 0.821552 | 1408404 |
| DM3770G_615 | 1675L | 0.879922 | 0.934937 | 0.822672 | 1408404 |
| DM3770G_615 | 991H | 0.878307 | 0.933908 | 0.820258 | 1408404 |
| DM3770G_615 | T0609 | 0.860142 | 0.935645 | 0.804788 | 1408404 |
| DM3770G_615 | T7901 | 0.90391 | 0.980828 | 0.88658 | 1408404 |
| DM3770G_615 | T7902 | 0.872341 | 0.932991 | 0.813886 | 1408404 |
| DM3770G_615 | T8402 | 0.904013 | 0.979725 | 0.885684 | 1408404 |
| DM3770G_615 | T8412 | 0.849266 | 0.932658 | 0.792075 | 1408404 |
| DM3770G_615 | T8415 | 0.908096 | 0.978877 | 0.888914 | 1408404 |
| DM3770G_615 | T8513 | 0.864238 | 0.933212 | 0.806517 | 1408404 |
| DM3770G_615 | T8602 | 0.915869 | 0.982445 | 0.899791 | 1408404 |
| DM3770G_994 | 2719K | 0.553608 | 0.964841 | 0.534144 | 677700 |
| KP2132G_2024 | 2141T | 0.951036 | 0.96525 | 0.917987 | 236499 |
| KP2132G_487 | 2141T | 0.751529 | 0.947831 | 0.712322 | 2396295 |
| KP2132G_543 | 2141T | 0.930654 | 0.947829 | 0.882101 | 7410314 |
| KP2132G_930 | 2141T | 0.913836 | 0.969131 | 0.885627 | 15633 |
| KP2133G_110 | 2141T | 0.970586 | 0.978075 | 0.949306 | 184775 |
| KP2133G_12 | 2141T | 0.852426 | 0.972543 | 0.829021 | 906223 |
| KP2133G_1401 | 2141T | 0.991723 | 0.918486 | 0.910884 | 36851 |
| KP2133G_1537 | 2141T | 0.937146 | 0.974138 | 0.91291 | 1388649 |
| KP2133G_1802 | 2141T | 0.885012 | 0.971645 | 0.859917 | 96236 |
| KP2133G_407 | 2141T | 0.514411 | 0.963889 | 0.495835 | 424581 |
| KP2133G_561 | 2141T | 0.847441 | 0.979638 | 0.830185 | 20805 |
| KP2133G_574 | 2141T | 0.956468 | 0.973746 | 0.931357 | 2478081 |
| KP2133G_581 | 2141T | 0.968286 | 0.976388 | 0.945423 | 52311 |
| KP2133G_742 | 2141T | 0.846229 | 0.972483 | 0.822943 | 860714 |
| KP3700G_1264 | 2141T | 0.916918 | 0.979339 | 0.897974 | 5185779 |
| KP3700G_1558 | 2141T | 0.715557 | 0.973669 | 0.696716 | 1902657 |
| KP3700G_2285 | 2141T | 0.523456 | 0.953913 | 0.499331 | 94260 |
| KP3700G_254 | 2141T | 0.65172 | 0.96095 | 0.62627 | 67506 |
| NR01_83_0.fasta | 1133Y | 0.67502 | 0.946155 | 0.638674 | 6971531 |
| NR01_83_0.fasta | 1675L | 0.6816 | 0.945046 | 0.644143 | 6971531 |
| NR01_83_0.fasta | 991H | 0.694159 | 0.946682 | 0.657148 | 6971531 |
| NR01_83_0.fasta | T0609 | 0.660568 | 0.946208 | 0.625035 | 6971531 |
| NR01_83_0.fasta | T7901 | 0.689513 | 0.945465 | 0.65191 | 6971531 |
| NR01_83_0.fasta | T7902 | 0.686353 | 0.946113 | 0.649367 | 6971531 |
| NR01_83_0.fasta | T8402 | 0.687244 | 0.944579 | 0.649156 | 6971531 |
| NR01_83_0.fasta | T8412 | 0.693179 | 0.945101 | 0.655124 | 6971531 |
| NR01_83_0.fasta | T8415 | 0.679432 | 0.944509 | 0.64173 | 6971531 |
| NR01_83_0.fasta | T8513 | 0.713256 | 0.953623 | 0.680177 | 6971531 |
| NR01_83_0.fasta | T8602 | 0.674679 | 0.944171 | 0.637012 | 6971531 |
| NR01_uc.fasta | 1133Y | 0.89634 | 0.937844 | 0.840627 | 1878871 |
| NR01_uc.fasta | 1675L | 0.900039 | 0.932247 | 0.839059 | 1878871 |
| NR01_uc.fasta | 991H | 0.896425 | 0.932008 | 0.835475 | 1878871 |
| NR01_uc.fasta | T0609 | 0.894596 | 0.932416 | 0.834136 | 1878871 |
| NR01_uc.fasta | T7901 | 0.908853 | 0.979432 | 0.89016 | 1878871 |
| NR01_uc.fasta | T7902 | 0.898114 | 0.931294 | 0.836408 | 1878871 |
| NR01_uc.fasta | T8402 | 0.912015 | 0.97896 | 0.892826 | 1878871 |
| NR01_uc.fasta | T8412 | 0.891515 | 0.931243 | 0.830217 | 1878871 |
| NR01_uc.fasta | T8415 | 0.912593 | 0.97762 | 0.892169 | 1878871 |
| NR01_uc.fasta | T8513 | 0.894387 | 0.932666 | 0.834164 | 1878871 |
| NR01_uc.fasta | T8602 | 0.915191 | 0.978577 | 0.895585 | 1878871 |
| NR02_1_1.fasta | 1133Y | 0.668416 | 0.931331 | 0.622517 | 6001350 |
| NR02_1_1.fasta | 1675L | 0.679296 | 0.925157 | 0.628455 | 6001350 |
| NR02_1_1.fasta | 991H | 0.683963 | 0.924842 | 0.632558 | 6001350 |
| NR02_1_1.fasta | T0609 | 0.658962 | 0.925538 | 0.609894 | 6001350 |
| NR02_1_1.fasta | T7901 | 0.692028 | 0.983431 | 0.680562 | 6001350 |
| NR02_1_1.fasta | T7902 | 0.674652 | 0.92368 | 0.623163 | 6001350 |
| NR02_1_1.fasta | T8402 | 0.69478 | 0.979676 | 0.680659 | 6001350 |
| NR02_1_1.fasta | T8412 | 0.675651 | 0.923138 | 0.623719 | 6001350 |
| NR02_1_1.fasta | T8415 | 0.69717 | 0.980238 | 0.683393 | 6001350 |
| NR02_1_1.fasta | T8513 | 0.682099 | 0.924662 | 0.630711 | 6001350 |
| NR02_1_1.fasta | T8602 | 0.67917 | 0.981851 | 0.666844 | 6001350 |
| NR03_1_5.fasta | 1133Y | 0.876318 | 0.936051 | 0.820278 | 62313 |
| NR03_1_5.fasta | 1675L | 0.910404 | 0.960303 | 0.874264 | 62313 |
| NR03_1_5.fasta | 991H | 0.932117 | 0.959799 | 0.894645 | 62313 |
| NR03_1_5.fasta | T0609 | 0.818304 | 0.954325 | 0.780928 | 62313 |
| NR03_1_5.fasta | T7901 | 0.955996 | 0.947273 | 0.905589 | 62313 |
| NR03_1_5.fasta | T7902 | 0.897549 | 0.955068 | 0.85722 | 62313 |
| NR03_1_5.fasta | T8402 | 0.880506 | 0.973973 | 0.857589 | 62313 |
| NR03_1_5.fasta | T8412 | 0.876318 | 0.940355 | 0.82405 | 62313 |
| NR03_1_5.fasta | T8415 | 0.880506 | 0.973992 | 0.857606 | 62313 |
| NR03_1_5.fasta | T8513 | 0.824804 | 0.947214 | 0.781266 | 62313 |
| NR03_1_5.fasta | T8602 | 0.862067 | 0.958841 | 0.826585 | 62313 |
| NR03_3_0.fasta | 1133Y | 0.668093 | 0.935554 | 0.625037 | 1713843 |
| NR03_3_0.fasta | 1675L | 0.681904 | 0.932979 | 0.636202 | 1713843 |
| NR03_3_0.fasta | 991H | 0.673016 | 0.932529 | 0.627607 | 1713843 |
| NR03_3_0.fasta | T0609 | 0.665706 | 0.93191 | 0.620378 | 1713843 |
| NR03_3_0.fasta | T7901 | 0.675478 | 0.965013 | 0.651845 | 1713843 |
| NR03_3_0.fasta | T7902 | 0.673639 | 0.932215 | 0.627976 | 1713843 |
| NR03_3_0.fasta | T8402 | 0.666042 | 0.962142 | 0.640827 | 1713843 |
| NR03_3_0.fasta | T8412 | 0.666094 | 0.931251 | 0.620301 | 1713843 |
| NR03_3_0.fasta | T8415 | 0.669292 | 0.963475 | 0.644846 | 1713843 |
| NR03_3_0.fasta | T8513 | 0.684772 | 0.933819 | 0.639453 | 1713843 |
| NR03_3_0.fasta | T8602 | 0.661184 | 0.963555 | 0.637087 | 1713843 |
| NR03_uc.fasta | 1133Y | 0.613876 | 0.946813 | 0.581226 | 7380431 |
| NR03_uc.fasta | 1675L | 0.620773 | 0.945993 | 0.587247 | 7380431 |
| NR03_uc.fasta | 991H | 0.638711 | 0.945861 | 0.604132 | 7380431 |
| NR03_uc.fasta | T0609 | 0.607121 | 0.945044 | 0.573756 | 7380431 |
| NR03_uc.fasta | T7901 | 0.622761 | 0.95804 | 0.59663 | 7380431 |
| NR03_uc.fasta | T7902 | 0.628333 | 0.945183 | 0.59389 | 7380431 |
| NR03_uc.fasta | T8402 | 0.638135 | 0.957549 | 0.611046 | 7380431 |
| NR03_uc.fasta | T8412 | 0.636491 | 0.943442 | 0.600492 | 7380431 |
| NR03_uc.fasta | T8415 | 0.628302 | 0.956369 | 0.600889 | 7380431 |
| NR03_uc.fasta | T8513 | 0.642384 | 0.946389 | 0.607945 | 7380431 |
| NR03_uc.fasta | T8602 | 0.615265 | 0.956948 | 0.588777 | 7380431 |
| T0609 | 1133Y | 0.885222 | 0.974391 | 0.862552 | 4392680 |
| T0609 | 1675L | 0.90846 | 0.978659 | 0.889073 | 4392680 |
| T0609 | 991H | 0.896712 | 0.982882 | 0.881362 | 4392680 |
| T0609 | NR01_83_0.fasta | 0.627597 | 0.93272 | 0.585372 | 4392680 |
| T0609 | NR02_1_1.fasta | 0.827633 | 0.920637 | 0.76195 | 4392680 |
| T0609 | NR03_uc.fasta | 0.636922 | 0.935509 | 0.595846 | 4392680 |
| T0609 | T7901 | 0.873915 | 0.924175 | 0.80765 | 4392680 |
| T0609 | T7902 | 0.894391 | 0.981676 | 0.878002 | 4392680 |
| T0609 | T8402 | 0.876731 | 0.926249 | 0.812071 | 4392680 |
| T0609 | T8412 | 0.895122 | 0.97822 | 0.875626 | 4392680 |
| T0609 | T8415 | 0.884963 | 0.923698 | 0.817439 | 4392680 |
| T0609 | T8513 | 0.892209 | 0.982359 | 0.87647 | 4392680 |
| T0609 | T8602 | 0.870858 | 0.923299 | 0.804062 | 4392680 |
| T0609 | TBF05_2_0.fasta | 0.809401 | 0.97434 | 0.788632 | 4392680 |
| T7901 | 1133Y | 0.889122 | 0.930659 | 0.827469 | 4485681 |
| T7901 | 1675L | 0.874732 | 0.924164 | 0.808396 | 4485681 |
| T7901 | 991H | 0.891699 | 0.923504 | 0.823488 | 4485681 |
| T7901 | NR01_83_0.fasta | 0.631319 | 0.946123 | 0.597305 | 4485681 |
| T7901 | NR02_1_1.fasta | 0.840667 | 0.979563 | 0.823486 | 4485681 |
| T7901 | NR03_uc.fasta | 0.626208 | 0.95685 | 0.599187 | 4485681 |
| T7901 | T0609 | 0.857654 | 0.924544 | 0.792939 | 4485681 |
| T7901 | T7902 | 0.8819 | 0.922027 | 0.813136 | 4485681 |
| T7901 | T8402 | 0.910335 | 0.981911 | 0.893868 | 4485681 |
| T7901 | T8412 | 0.880605 | 0.921929 | 0.811855 | 4485681 |
| T7901 | T8415 | 0.902292 | 0.980763 | 0.884935 | 4485681 |
| T7901 | T8513 | 0.87447 | 0.92361 | 0.807669 | 4485681 |
| T7901 | T8602 | 0.901004 | 0.982316 | 0.885071 | 4485681 |
| T7901 | TBF05_2_0.fasta | 0.796251 | 0.918111 | 0.731047 | 4485681 |
| T7902 | 1133Y | 0.846907 | 0.976243 | 0.826787 | 4679711 |
| T7902 | 1675L | 0.902333 | 0.985913 | 0.889622 | 4679711 |
| T7902 | 991H | 0.879235 | 0.982723 | 0.864044 | 4679711 |
| T7902 | NR01_83_0.fasta | 0.616936 | 0.927803 | 0.572395 | 4679711 |
| T7902 | NR02_1_1.fasta | 0.804973 | 0.917494 | 0.738558 | 4679711 |
| T7902 | NR03_uc.fasta | 0.625576 | 0.933852 | 0.584195 | 4679711 |
| T7902 | T0609 | 0.853667 | 0.979927 | 0.836531 | 4679711 |
| T7902 | T7901 | 0.853014 | 0.921915 | 0.786406 | 4679711 |
| T7902 | T8402 | 0.881667 | 0.921382 | 0.812352 | 4679711 |
| T7902 | T8412 | 0.887045 | 0.979357 | 0.868734 | 4679711 |
| T7902 | T8415 | 0.875878 | 0.921236 | 0.80689 | 4679711 |
| T7902 | T8513 | 0.882228 | 0.981673 | 0.866059 | 4679711 |
| T7902 | T8602 | 0.839603 | 0.922374 | 0.774428 | 4679711 |
| T7902 | TBF05_2_0.fasta | 0.770507 | 0.976281 | 0.752231 | 4679711 |
| T8402 | 1133Y | 0.880803 | 0.931718 | 0.82066 | 4501956 |
| T8402 | 1675L | 0.871153 | 0.925977 | 0.806668 | 4501956 |
| T8402 | 991H | 0.889551 | 0.925788 | 0.823536 | 4501956 |
| T8402 | NR01_83_0.fasta | 0.624453 | 0.944341 | 0.589697 | 4501956 |
| T8402 | NR02_1_1.fasta | 0.838445 | 0.976034 | 0.818351 | 4501956 |
| T8402 | NR03_uc.fasta | 0.640692 | 0.955863 | 0.612414 | 4501956 |
| T8402 | T0609 | 0.855565 | 0.926143 | 0.792376 | 4501956 |
| T8402 | T7901 | 0.911137 | 0.981586 | 0.894359 | 4501956 |
| T8402 | T7902 | 0.891047 | 0.9231 | 0.822525 | 4501956 |
| T8402 | T8412 | 0.891846 | 0.923396 | 0.823527 | 4501956 |
| T8402 | T8415 | 0.912321 | 0.979713 | 0.893813 | 4501956 |
| T8402 | T8513 | 0.879714 | 0.925564 | 0.814232 | 4501956 |
| T8402 | T8602 | 0.898003 | 0.980678 | 0.880652 | 4501956 |
| T8402 | TBF05_2_0.fasta | 0.788936 | 0.918738 | 0.724825 | 4501956 |
| T8412 | 1133Y | 0.890765 | 0.972143 | 0.865951 | 4467551 |
| T8412 | 1675L | 0.889963 | 0.979045 | 0.871314 | 4467551 |
| T8412 | 991H | 0.920565 | 0.98011 | 0.902255 | 4467551 |
| T8412 | NR01_83_0.fasta | 0.635071 | 0.930336 | 0.590829 | 4467551 |
| T8412 | NR02_1_1.fasta | 0.824026 | 0.916729 | 0.755409 | 4467551 |
| T8412 | NR03_uc.fasta | 0.638637 | 0.933147 | 0.595942 | 4467551 |
| T8412 | T0609 | 0.882977 | 0.978528 | 0.864018 | 4467551 |
| T8412 | T7901 | 0.891578 | 0.921872 | 0.821921 | 4467551 |
| T8412 | T7902 | 0.917881 | 0.979589 | 0.899146 | 4467551 |
| T8412 | T8402 | 0.901942 | 0.923388 | 0.832842 | 4467551 |
| T8412 | T8415 | 0.883512 | 0.921931 | 0.814537 | 4467551 |
| T8412 | T8513 | 0.918563 | 0.979996 | 0.900188 | 4467551 |
| T8412 | T8602 | 0.8739 | 0.922244 | 0.805949 | 4467551 |
| T8412 | TBF05_2_0.fasta | 0.816165 | 0.974634 | 0.795462 | 4467551 |
| T8415 | 1133Y | 0.87938 | 0.93364 | 0.821024 | 4476275 |
| T8415 | 1675L | 0.898434 | 0.926844 | 0.832708 | 4476275 |
| T8415 | 991H | 0.884153 | 0.925735 | 0.818491 | 4476275 |
| T8415 | NR01_83_0.fasta | 0.636397 | 0.946789 | 0.602534 | 4476275 |
| T8415 | NR02_1_1.fasta | 0.852261 | 0.977522 | 0.833104 | 4476275 |
| T8415 | NR03_uc.fasta | 0.641977 | 0.956194 | 0.613855 | 4476275 |
| T8415 | T0609 | 0.86685 | 0.924166 | 0.801113 | 4476275 |
| T8415 | T7901 | 0.906795 | 0.981376 | 0.889907 | 4476275 |
| T8415 | T7902 | 0.884674 | 0.924949 | 0.818278 | 4476275 |
| T8415 | T8402 | 0.91731 | 0.981868 | 0.900677 | 4476275 |
| T8415 | T8412 | 0.877572 | 0.924005 | 0.810881 | 4476275 |
| T8415 | T8513 | 0.876244 | 0.925177 | 0.810681 | 4476275 |
| T8415 | T8602 | 0.889734 | 0.982798 | 0.874429 | 4476275 |
| T8415 | TBF05_2_0.fasta | 0.779922 | 0.91913 | 0.71685 | 4476275 |
| T8513 | 1133Y | 0.882059 | 0.974339 | 0.859424 | 4559547 |
| T8513 | 1675L | 0.889501 | 0.981425 | 0.872979 | 4559547 |
| T8513 | 991H | 0.894236 | 0.983704 | 0.879664 | 4559547 |
| T8513 | NR01_83_0.fasta | 0.649385 | 0.938047 | 0.609154 | 4559547 |
| T8513 | NR02_1_1.fasta | 0.817981 | 0.917444 | 0.750452 | 4559547 |
| T8513 | NR03_uc.fasta | 0.635693 | 0.936753 | 0.595487 | 4559547 |
| T8513 | T0609 | 0.861942 | 0.982137 | 0.846545 | 4559547 |
| T8513 | T7901 | 0.870331 | 0.92317 | 0.803463 | 4559547 |
| T8513 | T7902 | 0.895824 | 0.981855 | 0.879569 | 4559547 |
| T8513 | T8402 | 0.873841 | 0.925228 | 0.808502 | 4559547 |
| T8513 | T8412 | 0.901188 | 0.980463 | 0.883581 | 4559547 |
| T8513 | T8415 | 0.863506 | 0.924296 | 0.798135 | 4559547 |
| T8513 | T8602 | 0.862103 | 0.920981 | 0.79398 | 4559547 |
| T8513 | TBF05_2_0.fasta | 0.798762 | 0.975963 | 0.779562 | 4559547 |
| T8602 | 1133Y | 0.892922 | 0.931257 | 0.83154 | 4444880 |
| T8602 | 1675L | 0.866881 | 0.924539 | 0.801465 | 4444880 |
| T8602 | 991H | 0.885145 | 0.922655 | 0.816683 | 4444880 |
| T8602 | NR01_83_0.fasta | 0.62017 | 0.945379 | 0.586296 | 4444880 |
| T8602 | NR02_1_1.fasta | 0.835341 | 0.977696 | 0.81671 | 4444880 |
| T8602 | NR03_uc.fasta | 0.626023 | 0.955289 | 0.598033 | 4444880 |
| T8602 | T0609 | 0.862304 | 0.923203 | 0.796082 | 4444880 |
| T8602 | T7901 | 0.909376 | 0.982195 | 0.893185 | 4444880 |
| T8602 | T7902 | 0.87942 | 0.922876 | 0.811596 | 4444880 |
| T8602 | T8402 | 0.910518 | 0.9807 | 0.892945 | 4444880 |
| T8602 | T8412 | 0.877947 | 0.922503 | 0.809909 | 4444880 |
| T8602 | T8415 | 0.896389 | 0.981936 | 0.880197 | 4444880 |
| T8602 | T8513 | 0.881462 | 0.921479 | 0.812249 | 4444880 |
| T8602 | TBF05_2_0.fasta | 0.80329 | 0.917094 | 0.736692 | 4444880 |
| TBF02_1_1.fasta | 1162T | 0.960455 | 0.984263 | 0.94534 | 450549 |
| TBF02_3_0.fasta | 1133Y | 0.8044 | 0.930773 | 0.748714 | 3234669 |
| TBF02_3_0.fasta | 1675L | 0.788952 | 0.924999 | 0.72978 | 3234669 |
| TBF02_3_0.fasta | 991H | 0.808884 | 0.924883 | 0.748123 | 3234669 |
| TBF02_3_0.fasta | T0609 | 0.783988 | 0.925451 | 0.725542 | 3234669 |
| TBF02_3_0.fasta | T7901 | 0.820008 | 0.976303 | 0.800576 | 3234669 |
| TBF02_3_0.fasta | T7902 | 0.794959 | 0.924469 | 0.734915 | 3234669 |
| TBF02_3_0.fasta | T8402 | 0.813269 | 0.975412 | 0.793272 | 3234669 |
| TBF02_3_0.fasta | T8412 | 0.802364 | 0.923339 | 0.740854 | 3234669 |
| TBF02_3_0.fasta | T8415 | 0.798753 | 0.975965 | 0.779555 | 3234669 |
| TBF02_3_0.fasta | T8513 | 0.798417 | 0.922979 | 0.736922 | 3234669 |
| TBF02_3_0.fasta | T8602 | 0.813756 | 0.978318 | 0.796112 | 3234669 |
| TBF03_2_0.fasta | 1133Y | 0.77877 | 0.972509 | 0.757361 | 4012364 |
| TBF03_2_0.fasta | 1675L | 0.769773 | 0.97976 | 0.754193 | 4012364 |
| TBF03_2_0.fasta | 991H | 0.792989 | 0.980324 | 0.777386 | 4012364 |
| TBF03_2_0.fasta | T0609 | 0.75793 | 0.979196 | 0.742162 | 4012364 |
| TBF03_2_0.fasta | T7901 | 0.77334 | 0.919818 | 0.711332 | 4012364 |
| TBF03_2_0.fasta | T7902 | 0.784174 | 0.980219 | 0.768662 | 4012364 |
| TBF03_2_0.fasta | T8402 | 0.768395 | 0.921958 | 0.708428 | 4012364 |
| TBF03_2_0.fasta | T8412 | 0.785248 | 0.97951 | 0.769158 | 4012364 |
| TBF03_2_0.fasta | T8415 | 0.752277 | 0.921598 | 0.693297 | 4012364 |
| TBF03_2_0.fasta | T8513 | 0.781884 | 0.981158 | 0.767152 | 4012364 |
| TBF03_2_0.fasta | T8602 | 0.75842 | 0.919919 | 0.697685 | 4012364 |
| TBF05_2_0.fasta | 1133Y | 0.92992 | 0.973602 | 0.905372 | 4056823 |
| TBF05_2_0.fasta | 1675L | 0.91262 | 0.980719 | 0.895024 | 4056823 |
| TBF05_2_0.fasta | 991H | 0.935601 | 0.981484 | 0.918277 | 4056823 |
| TBF05_2_0.fasta | T0609 | 0.915587 | 0.980034 | 0.897306 | 4056823 |
| TBF05_2_0.fasta | T7901 | 0.920769 | 0.92287 | 0.84975 | 4056823 |
| TBF05_2_0.fasta | T7902 | 0.923265 | 0.981124 | 0.905837 | 4056823 |
| TBF05_2_0.fasta | T8402 | 0.912109 | 0.923725 | 0.842538 | 4056823 |
| TBF05_2_0.fasta | T8412 | 0.933926 | 0.98047 | 0.915686 | 4056823 |
| TBF05_2_0.fasta | T8415 | 0.90229 | 0.923779 | 0.833517 | 4056823 |
| TBF05_2_0.fasta | T8513 | 0.932749 | 0.982141 | 0.916091 | 4056823 |
| TBF05_2_0.fasta | T8602 | 0.918734 | 0.921887 | 0.846969 | 4056823 |
| TBF05_uc.fasta | 1133Y | 0.659554 | 0.965507 | 0.636804 | 198111 |
| TBF05_uc.fasta | 1675L | 0.646047 | 0.977428 | 0.631464 | 198111 |
| TBF05_uc.fasta | 991H | 0.640762 | 0.978234 | 0.626815 | 198111 |
| TBF05_uc.fasta | T0609 | 0.667929 | 0.972741 | 0.649722 | 198111 |
| TBF05_uc.fasta | T7901 | 0.634311 | 0.91599 | 0.581023 | 198111 |
| TBF05_uc.fasta | T7902 | 0.630964 | 0.977544 | 0.616795 | 198111 |
| TBF05_uc.fasta | T8402 | 0.668655 | 0.919369 | 0.614741 | 198111 |
| TBF05_uc.fasta | T8412 | 0.665173 | 0.966466 | 0.642867 | 198111 |
| TBF05_uc.fasta | T8415 | 0.635492 | 0.916043 | 0.582138 | 198111 |
| TBF05_uc.fasta | T8513 | 0.631192 | 0.97748 | 0.616978 | 198111 |
| TBF05_uc.fasta | T8602 | 0.644336 | 0.912996 | 0.588276 | 198111 |
| TBF07_1_1.fasta | 1162T | 0.895614 | 0.984754 | 0.881959 | 743952 |
| TBF09_17_0.fasta | 1133Y | 0.953758 | 0.971264 | 0.926351 | 1861029 |
| TBF09_17_0.fasta | 1675L | 0.939717 | 0.978883 | 0.919873 | 1861029 |
| TBF09_17_0.fasta | 991H | 0.966973 | 0.979451 | 0.947103 | 1861029 |
| TBF09_17_0.fasta | T0609 | 0.930403 | 0.97855 | 0.910446 | 1861029 |
| TBF09_17_0.fasta | T7901 | 0.943027 | 0.922276 | 0.869731 | 1861029 |
| TBF09_17_0.fasta | T7902 | 0.956445 | 0.979964 | 0.937282 | 1861029 |
| TBF09_17_0.fasta | T8402 | 0.937665 | 0.924468 | 0.866841 | 1861029 |
| TBF09_17_0.fasta | T8412 | 0.96116 | 0.979005 | 0.94098 | 1861029 |
| TBF09_17_0.fasta | T8415 | 0.920284 | 0.923824 | 0.85018 | 1861029 |
| TBF09_17_0.fasta | T8513 | 0.955093 | 0.979827 | 0.935826 | 1861029 |
| TBF09_17_0.fasta | T8602 | 0.934865 | 0.921081 | 0.861086 | 1861029 |
| TBF09_2_0.fasta | 1162T | 0.943394 | 0.981959 | 0.926374 | 294653 |

**Table S3.**
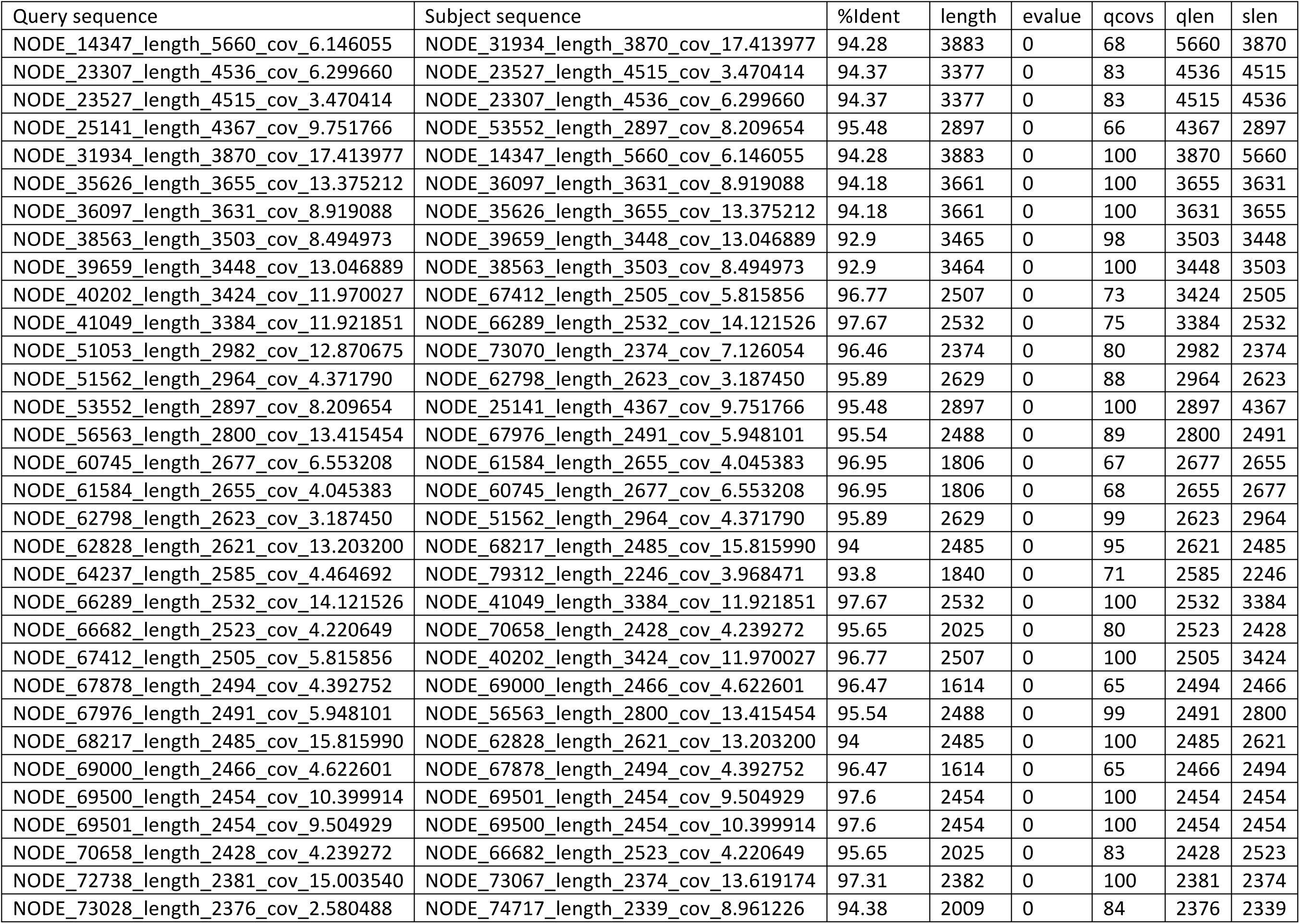
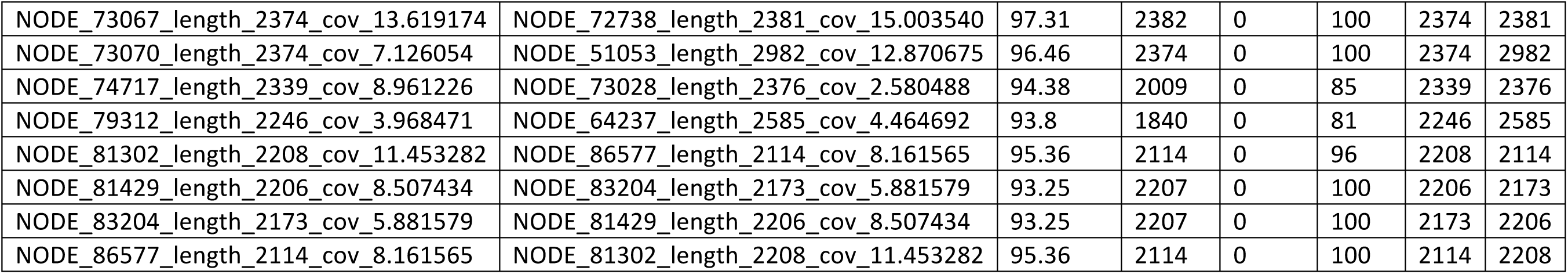
Comparison of contigs in the BSC2 bin.

**Table S4.** List of GCFs found in this study.

|  |  |  |  |
| --- | --- | --- | --- |
| **GCF_1** | **cf_fatty_acid-t1pks-nrps** | **GCF_62** | **terpene** |
| **GCF_2** | bacteriocin-transatpks-t1pks-nrps | **GCF_63** | terpene |
| **GCF_3** | cf_fatty_acid-transatpks-t1pks-nrps | **GCF_64** | terpene |
| **GCF_4** | t1pks-cf_saccharide-nrps | **GCF_65** | terpene |
| **GCF_5** | terpene-arylpolyene | **GCF_66** | t1pks |
| **GCF_6** | transatpks-cf_saccharide-nrps | **GCF_67** | t1pks |
| **GCF_7** | nrps | **GCF_68** | t1pks |
| **GCF_8** | cf_fatty_acid-nrps_(tunerbactin) | **GCF_69** | t1pks |
| **GCF_9** | t1pks | **GCF_70** | t1pks-PUFA |
| **GCF_10** | hserlactone-transatpks-nrps | **GCF_71** | t1pks-nrps |
| **GCF_11** | transatpks_(tartrolon) | **GCF_72** | t1pks-nrps |
| **GCF_12** | transatpks-nrps | **GCF_73** | t1pks-nrps |
| **GCF_13** | t1pks-nrps | **GCF_74** | t1pks-nrps |
| **GCF_14** | siderophore | **GCF_75** | t1pks-nrps |
| **GCF_15** | transatpks-otherks | **GCF_76** | t1pks-cf_saccharide-nrps |
| **GCF_16** | t1pks-nrps | **GCF_77** | t1pks-cf_saccharide-nrps |
| **GCF_17** | nrps | **GCF_78** | t1pks-cf_fatty_acid |
| **GCF_18** | transatpks | **GCF_79** | siderophore |
| **GCF_19** | t1pks | **GCF_80** | nrps-transatpks-otherks |
| **GCF_20** | nrps | **GCF_81** | nrps |
| **GCF_21** | transatpks | **GCF_82** | nrps |
| **GCF_22** | t1pks | **GCF_83** | nrps |
| **GCF_23** | siderophore | **GCF_84** | nrps |
| **GCF_24** | nrps | **GCF_85** | nrps |
| **GCF_25** | nrps | **GCF_86** | nrps |
| **GCF_26** | transatpks | **GCF_87** | nrps |
| **GCF_27** | transatpks-otherks-nrps | **GCF_88** | nrps |
| **GCF_28** | nrps | **GCF_89** | nrps |
| **GCF_29** | nrps | **GCF_90** | nrps |
| **GCF_30** | nrps | **GCF_91** | nrps |
| **GCF_31** | nrps | **GCF_92** | nrps |
| **GCF_32** | transatpks | **GCF_93** | nrps |
| **GCF_33** | transatpks-t1pks-nrps | **GCF_94** | nrps |
| **GCF_34** | thiopeptide-<br>hserlactone | **GCF_95** | nrps |
| **GCF_35** | t1pks-cf_saccharide-nrps | **GCF_96** | nrps |
| **GCF_36** | t1pks-nrps | **GCF_97** | nrps |
| **GCF_37** | t1pks-nrps | **GCF_98** | nrps |
| **GCF_38** | t1pks-cf_saccharide-nrps | **GCF_99** | nrps |
| **GCF_39** | nrps | **GCF_100** | nrps |
| **GCF_40** | nrps | **GCF_101** | nrps |
| **GCF_41** | nrps | **GCF_102** | nrps |
| **GCF_42** | nrps | **GCF_103** | nrps |
| **GCF_43** | nrps | **GCF_104** | nrps |
| **GCF_44** | nrps | **GCF_105** | nrps |
| **GCF_45** | nrps | **GCF_106** | nrps |
| **GCF_46** | nrps | **GCF_107** | nrps |
| **GCF_47** | nrps | **GCF_108** | nrps |
| **GCF_48** | nrps | **GCF_109** | nrps |
| **GCF_49** | nrps | **GCF_110** | nrps |
| **GCF_50** | nrps | **GCF_111** | nrps |
| **GCF_51** | hserlactone-t1pks-nrps | **GCF_112** | nrps |
| **GCF_52** | cf_saccharide-nrps | **GCF_113** | nrps |
| **GCF_53** | transatpks | **GCF_114** | nrps |
| **GCF_54** | transatpks | **GCF_115** | nrps |
| **GCF_55** | transatpks | **GCF_116** | nrps |
| **GCF_56** | transatpks | **GCF_117** | hserlactone-transatpks-cf_fatty_acid |
| **GCF_57** | transatpks | **GCF_118** | hserlactone-nrps |
| **GCF_58** | transatpks-t1pks-nrps | **GCF_119** | cf_saccharide-nrps |
| **GCF_59** | transatpks-otherks | **GCF_120** | cf_fatty_acid-t1pks |
| **GCF_60** | transatpks-cf_saccharide | **GCF_121** | bacteriocin-<br>lantipeptide |
| **GCF_61** | transatpks-cf_fatty_acid | **GCF_122** | arylpolyene-<br>nrps_(butunamide) |

